# CHOP promotes the transition to chronic integrated stress response signaling with suppression of hepatocyte identity

**DOI:** 10.64898/2026.05.13.724984

**Authors:** Theo F. Velarde, Kaihua Liu, Zewei Zhang, Reed C. Adajar, Chaoxian Zhao, Huojun Cao, D. Thomas Rutkowski

## Abstract

The transcription factor CHOP promotes cell death during ER stress, but it is strongly induced even by moderate stresses that do not result in appreciable cell death. Its role during less severe stresses—especially in intact tissues *in vivo*—is poorly understood. Here, we both deleted and restored CHOP specifically in hepatocytes and challenged animals with ER stress *in vivo*. We found that CHOP influenced stress-dependent hepatocyte gene expression through two previously unappreciated mechanisms. It directly suppressed the expression of transcriptional master regulators of hepatocyte identity and metabolism. And more broadly, it exacerbated ER stress through the promotion of protein synthesis, which led to persistent activation of the integrated stress response (ISR) despite dephosphorylation of eIF2α. This shift to second-phase ISR signaling was phenocopied by deletion of the protective UPR sensor ATF6α, suggesting that it reflects a transition from an acute stress response to a chronic one. Our findings show that CHOP augments the capacity of the ISR and UPR to continue to mount a protective response even after eIF2α phosphorylation has been suppressed. *In vivo*, where ISR signaling intersects with hepatocyte gene regulatory networks, this transition favors lipid dysregulation, highlighting a pathway through which CHOP impacts tissue function independent of cell death.

## Introduction

Cellular stress responses act through both transcriptional and non-transcriptional mechanisms to restore homeostasis. For a diverse group of stressors that tax the cellular protein synthetic and folding machinery—including ER stress, viral infection, amino acid deprivation, and heme depletion among others—a major early response to stress is phosphorylation of the translation initiation factor eIF2α, which initiates the integrated stress response (ISR) (Costa-Mattioli and Walter 2020; Harding et al. 2003). Although eIF2α phosphorylation rapidly inhibits global protein synthesis, it must necessarily be transient to allow cellular recovery. Yet many cellular stresses persist beyond the window during which eIF2α is phosphorylated. How dephosphorylation of eIF2α and the resumption of protein synthesis affect the ability of cells to withstand longer-term insults is not well understood.

PERK (protein kinase R-like endoplasmic reticulum kinase) catalyzes eIF2α phosphorylation during ER stress, inhibiting global protein synthesis while launching the ISR that exerts broad control over the transcriptome (Sood et al. 2000; Harding et al. 2000). Acting in concert are the IRE1α (Inositol-requiring 1) and ATF6α (activating transcription factor 6) pathways, the three of which together comprise the unfolded protein response (UPR) (Hetz, Zhang, and Kaufman 2020). The relationships among the three pathways are complex. Although IRE1α is the most conserved of these and both IRE1α and ATF6α are uniquely required for cells to adequately withstand various developmental and pathological challenges, during conventional ER stresses the large majority of UPR-dependent gene regulation appears to require PERK at least where the contributions of each pathway have been systematically studied (Adamson et al. 2016; Teske et al. 2011; Wu et al. 2007). Therefore, these pathways are responsive to homeostasis in the ER both directly, and indirectly through PERK/eIF2α signaling.

The CHOP (C/EBP-homologous) protein is of particular consequence for UPR signaling and restoration of homeostasis. During severe ER stresses of the sort typically studied experimentally, CHOP promotes stress-induced cell death (Yang et al. 2017; Zinszner et al. 1998). Its deletion *in vivo* typically diminishes cell death in mouse models of pathology, while also often ameliorating disease phenotypes. However, whether CHOP-induced cell death causes disease or only correlates with it has not generally been tested, and in some cases its effects on cell death appear separable from its effects on disease (Ji et al. 2005; Pennuto et al. 2008). Thus, how CHOP exacerbates disease, be it through cell death or other mechanisms, is not well understood.

CHOP does not appear to function as a cell death regulator *per se*. It is a member of the C/EBP family of transcription factors, which regulate cell proliferation and differentiation; CHOP has an altered DNA binding capacity (Barone et al. 1994) and a truncated and disordered N-terminus (Singh et al. 2008; Canales et al. 2017) relative to the other members of the family, which are themselves part of the larger category of transcription factors known as bZIPs (basic leucine zipper) that form homo- and hetero-dimers. Although it has been proposed to transcriptionally regulate various pro- and anti-apoptotic genes (Puthalakath et al. 2007; Yamaguchi and Wang 2004; Galehdar et al. 2010), at the whole genome level its best characterized effect is the stimulation of protein synthesis. It carries out this role by transcriptionally coregulating with the ISR-regulated factor ATF4 not only genes encoding various tRNA synthetases, ribosomal components, and translation regulatory factors, but also by enhancing expression of the eIF2α phosphatase GADD34 (Han et al. 2013; Krokowski et al. 2013; Marciniak et al. 2004). During severe ER stress, it is likely that the promotion of protein synthesis oxidation tax an already overburdened organelle, accelerating the cytotoxic consequences of ER disruption including oxidative damage (Li et al. 2009; Harding et al. 2003). However, CHOP is robustly induced even during mild ER stresses that do not induce significant cell death (Rutkowski et al. 2006). The impact of CHOP’s regulation of ISR signaling during ongoing exposure to non-lethal stress is largely unknown.

Using a newly created multifunctional targeted allele of the *Chop* gene, we recently showed that, within a population of cultured MEFs (mouse embryonic fibroblasts), CHOP promotes both cell death and, paradoxically, adaptation and proliferation (Liu et al. 2024). In that context, the resumption of protein synthesis exacerbated ER stress but also maximized the activation of the UPR transcriptional program, thereby promoting restoration of homeostasis and subsequent deactivation of the UPR. Thus, we proposed that CHOP facilitates adaptation by stress testing the ER.

Here, we turned our attention to the role of CHOP in an intact tissue *in vivo*, examining its contribution to ER stress signaling in the liver. We have previously shown that, in the liver in response to stress caused by either inhibition of N-glycosylation or proteasome inhibition, CHOP contributes to the direct regulation of several genes encoding transcriptional master regulators of metabolism (Chikka et al. 2013). This function differentiates the contribution of CHOP in the liver UPR from that in MEFs. Whole-body deletion of CHOP protects against several forms of liver injury (DeZwaan-McCabe et al. 2013; Scaiewicz et al. 2013; Zhou et al. 2023; Yang et al. 2022; Uzi et al. 2013), underscoring the importance of determining its contribution to ER stress signaling in hepatocytes in particular. By manipulating *Chop* specifically in hepatocytes, we show here that CHOP promotes the transition of the ISR from acute to chronic signaling characterized by maintenance of the ATF4-dependent transcriptional program despite eIF2α dephosphorylation, and suppression of genes involved in hepatocyte metabolism and identity.

## Results

### CHOP amplifies hepatic steatosis and ISR signaling despite eIF2α dephosphorylation

The ER stress-inducing agent tunicamycin (TM) is widely used to elicit ER stress *in vivo*. Injected intraperitoneally, it efficiently inhibits N-linked protein glycosylation and thereby elicits ER stress in the liver and, to a lesser extent, the kidneys (Zinszner et al. 1998; DeZwaan-McCabe et al. 2017). Because of its *in vivo* potency and because N-glycosylation is an ER-specific post-translational modification, TM challenge is a robust method for examining the hepatic response to ER stress. Its most visible impact on the liver is mild lipid accumulation, or steatosis, that occurs in both male and female mice (Fig. 1A). In whole livers from animals of both sexes, expression of CHOP uniformly peaks 8 hours after challenge and diminishes thereafter, although the rate of decline in CHOP expression varied somewhat among animals (Fig. 1B). Consistent with intraperitoneally delivered TM being taken up through the portal circulation, expression of CHOP was higher in periportal and intermediate regions than in the pericentral region. Its staining pattern was predominantly nuclear (Fig. S1). The fact that CHOP is most abundant 8 hours after challenge makes this the most logical time point for understanding its role in the liver. Although the best described consequence of CHOP expression is cell death, the typical experimental dose of TM induces death in only a trivial fraction of cells, even at later time points (Fig. 1C). Therefore, the effects of CHOP on liver physiology under these conditions are not a consequence of cell death.

**Figure 1:**
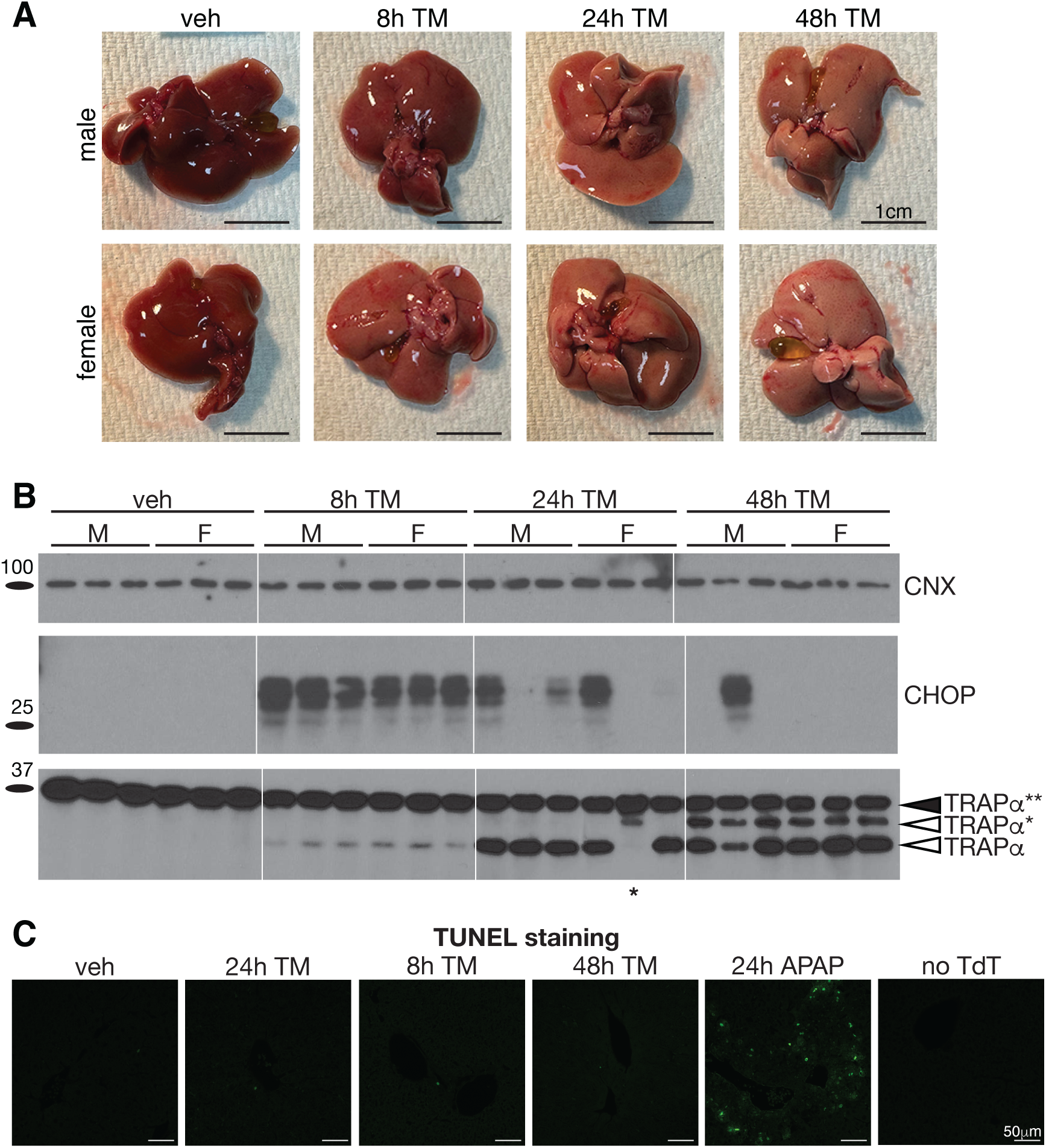
Tunicamycin challenge elicits ER stress and CHOP induction in male and female mice without significant cell death **(A)** Whole livers from male or female mice challenged for the indicated times with 1 mg/kg b.w. TM or vehicle (veh) **(B)** Immunoblot to detect CHOP and the ER-resident glycoprotein TRAPα, which exists normally as a doubly glycosylated form (TRAPα**) and for which mono-glycosylated (TRAPα*) and unglycosylated (TRAPα) forms are induced by TM. Calnexin (CNX) was used as a loading control. Asterisk under lane 17 designates an animal that did not receive a fully effective dose of TM, as indicated by very modest inhibition of TRAPα glycosylation. Here and elsewhere, hairlines are used only for visual clarity. **(C)** TUNEL staining was used to detect cell death at the indicated time points, with 300 mg/kg acetaminophen for 24h used as a positive control.

We have previously reported the creation of a novel “knockout-first” targeted allele of CHOP (*Chop^FLuL/FLuL^*) in which CHOP expression can be restored by FLP-mediated excision of a gene trap cassette (*Chop^fl/fl^*) and thereafter knocked out by CRE-mediated excision of the entire CHOP ORF (*Chop^KO^*) (Fig. 2A) (Liu et al. 2024). *Floxed* animals were bred into the nTnG CRE indicator line in which CRE action converts nuclear fluorescence from red (tdTomato) to green (EGFP) (Prigge et al. 2013). We used AAV-TBG-CRE to delete *Chop* specifically in hepatocytes of these adult animals (*Chop^HKO^*), with AAV-TBG-GFP serving as a control (Kiourtis et al. 2021). In the former group, the large majority of hepatocytes would be expected to be CHOP-deleted with green nuclear fluorescence, while non-hepatocytes would be CHOP-competent with red nuclear fluorescence. In AAV-TBG-GFP-injected animals, most hepatocytes would be green in the cytoplasm and red in the nucleus, whereas non-hepatocytes would remain red in the nucleus.

**Figure 2:**
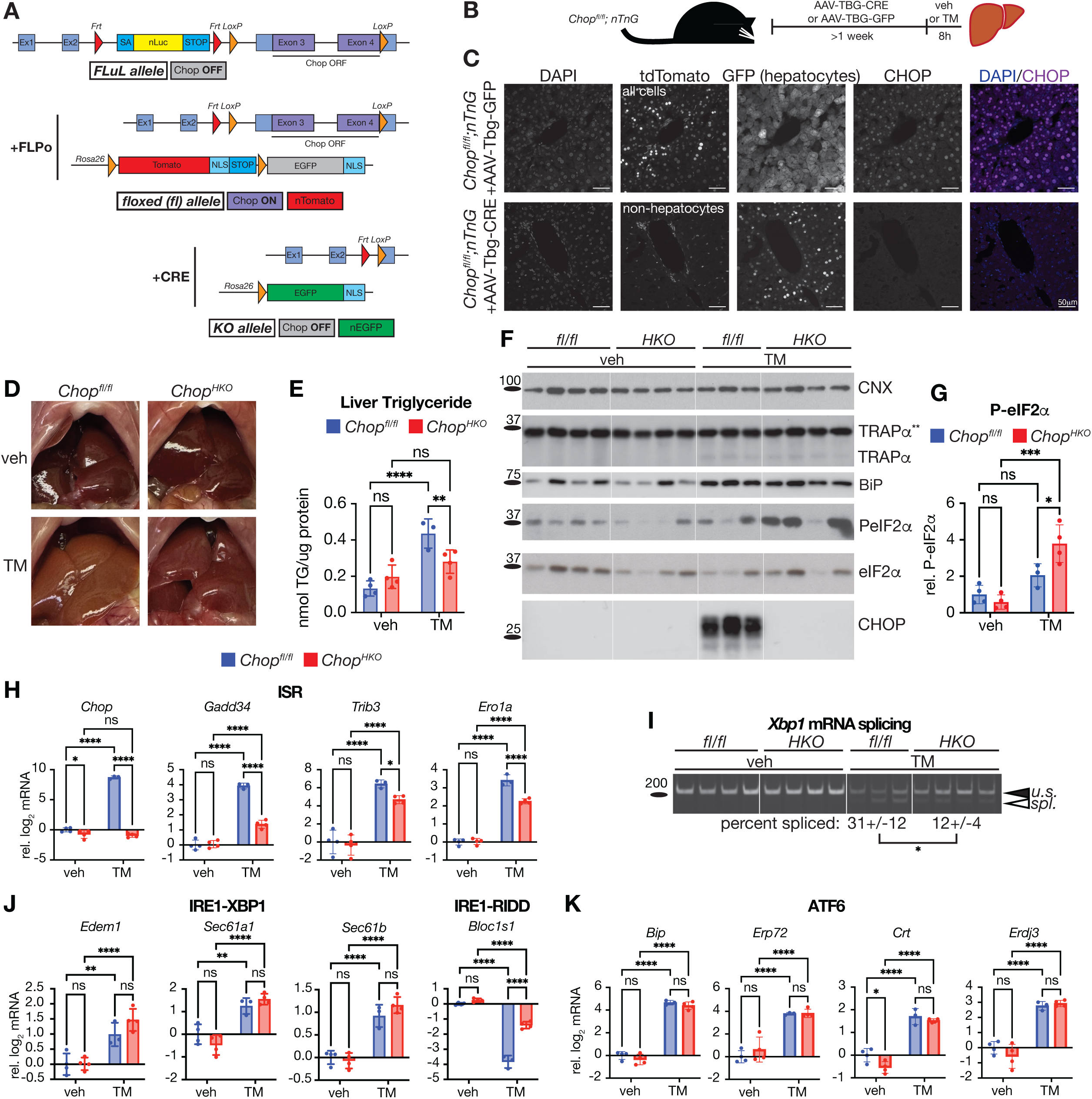
Hepatocyte-specific deletion of CHOP attenuates IRE1 activation and expression of ISR-regulated genes, but not XBP1- or ATF6-regulated genes **(A)** Schematic showing the *Chop FLuL* allele, in which CHOP expression is silenced by a gene trap; the *floxed* allele, in which CHOP expression is wild-type by virtue of FLPo-mediated excision of the gene trap; and the deleted allele, in which the entire open reading frame of CHOP is deleted by CRE. Also shown is the nTnG CRE reporter allele bred into *Chop^fl/fl^* mice, which converts from expression of tdTomato to EGFP upon CRE action. Figure adapted from (Liu et al. 2024). **(B)** Experimental paradigm for *Chop* deletion in *Chop^fl/fl^*; *nTnG* animals and challenge with 1 mg/kg TM **(C)** tdTomato, EGFP, and nuclei (DAPI) were visualized, and CHOP detected by immunostaining, in livers of an animal with an intact *Chop* allele expressing nuclear tdTomato in all cells and cytoplasmic AAV-driven EGFP in hepatocytes (top row), and an animal with a deleted *Chop* allele expressing nuclear tdTomato in all non-hepatocytes and nuclear EGFP in hepatocytes (bottom row). Note the absence of CHOP expression in the liver when CHOP is deleted specifically in hepatocytes. **(D)** *In situ* liver images from non-deleted (*fl/fl*) or hepatocyte-deleted (*HKO*) animals after challenge **(E)** Direct measurement of liver triglyceride normalized against protein content. **(F)** Immunoblot showing inhibition of glycosylation by TM using TRAPα as an indicator, as well as expression of the ER chaperone and UPR target gene BiP and phosphorylated and total species of eIF2α. Consistent with Figure 1, CHOP itself is undetectable in unchallenged livers. **(G)** Quantification of eIF2α phosphorylation from (F) **(H)** qRT-PCR detection of the indicated ISR target genes **(I)** *Xbp1* mRNA splicing detected by conventional RT-PCR with primers that equivalently recognize both spliced and unspliced forms **(J, K)** qRT-PCR detection of targets of the IRE1-XBP1 axis (J), a canonical RIDD target (*Bloc1s1*; J), and targets of the ATF6 axis (K). Here and elsewhere, error bars represent means +/- SDM and statistical comparisons were by two-way ANOVA.

After deletion, animals were challenged with TM for 8 hours (Fig. 2B), revealing that only hepatocytes express CHOP in response to TM. This conclusion was supported by the fact that all CHOP-expressing nuclei in AAV-TBG-GFP-expressing animals showed green cytoplasmic staining, and the fact that CRE-mediated deletion of CHOP specifically in hepatocytes abolished CHOP immunodetection in the liver (Fig. 2C). That CHOP expression is restricted to hepatocytes is consistent with the observation that the putative TM transporter MFSD2A is not expressed in non-hepatocyte liver cell types (Pu et al. 2016).

Deletion of CHOP had no apparent effect on the liver absent an ER stress. In contrast, a TM challenge produced more pronounced hepatic steatosis in wild-type mice than in animals with a hepatocyte-specific deletion of CHOP (*Chop^HKO^*), seen by the hue of whole liver (Fig. 2D) and by direct measurement of hepatic triglyceride (Fig. 2E). This finding confirms that CHOP influences hepatocyte metabolism during an ER stress challenge.

Focused examination of signaling from UPR pathways revealed a discordance between the status of the upstream UPR sensors and well-characterized downstream target genes (Adamson et al. 2016). CHOP is well-known to transcriptionally induce expression of the eIF2α phosphatase subunit GADD34 (Marciniak et al. 2004). This regulation has been postulated to both promote cell death during severe stress and adaptation during mild stress (Marciniak et al. 2004; Liu et al. 2024). As expected, eIF2α phosphorylation was elevated in TM-challenged *Chop^HKO^* animals compared to non-deleted animals (Fig. 2F, G). However, expression of ATF4 was not elevated in *Chop^HKO^*livers; if anything, it was suppressed (Fig. S2A). The expression of representative ISR target genes was diminished by CHOP deletion (Fig. 2H). In contrast, CHOP appeared to enhance activation of IRE1, seen by elevated splicing of the IRE1 endoribonuclease target *Xbp1* in non-deleted animals (Fig. 2I). Yet XBP1-dependent genes were not significantly affected by loss of CHOP (Fig. 2J), nor were genes regulated by the ATF6α pathway (Fig. 2K). Strikingly, expression of the gene *Bloc1s1* was much more strongly suppressed in CHOP-expressing livers than in *Chop^HKO^* livers (Fig. 2J). *Bloc1s1* is a canonical target of regulated IRE1-dependent decay (RIDD) (Bright et al. 2015; Hollien et al. 2009), and both its suppression and the augmented splicing of *Xbp1* in CHOP-expressing livers suggests that CHOP enhances IRE1 activation but without a concomitant increase in XBP1-dependent transcription. The enhancement of IRE1 activation is most likely due to enhanced ER stress caused by the resumption of protein synthesis, because treatment of primary hepatocytes *ex vivo* with the ISR inhibitor ISRIB—which prevents the suppression of protein synthesis caused by eIF2α phosphorylation (Sidrauski et al. 2013)—had the same effects on *Xbp1* splicing and RIDD as did CHOP (Fig. S2B, C).

### CHOP broadly enhances ISR-dependent gene regulation

We used RNA-seq for an unbiased assessment of CHOP’s effects on hepatic gene expression (Worksheet S1). For this experiment we used *Chop^FLuL/FLuL^*animals injected with either AAV-TBG-GFP or AAV-TBG-FLPo. *Chop^FLuL/FLuL^*animals are incompetent to express CHOP in all cells, whereas AAV-TBG-FLPo-treatment restores CHOP competence specifically in hepatocytes (Fig. 2A). Both male and female animals were used, and principal component analysis suggested that, in both sexes, CHOP restoration had little effect on gene expression in the absence of a stress challenge (Fig. 3A). Conversely, during stress treatment, livers lacking CHOP were shifted to a profile closer to untreated animals than livers expressing CHOP. Globally, this finding suggests that CHOP broadly enhances UPR signaling. Supporting this idea, splicing of *Xbp1* mRNA was more robust in CHOP-expressing livers, similarly to *Chop^fl/fl^* vs. *Chop^LKO^*livers (Fig. 3B and Fig. 2I).

**Figure 3:**
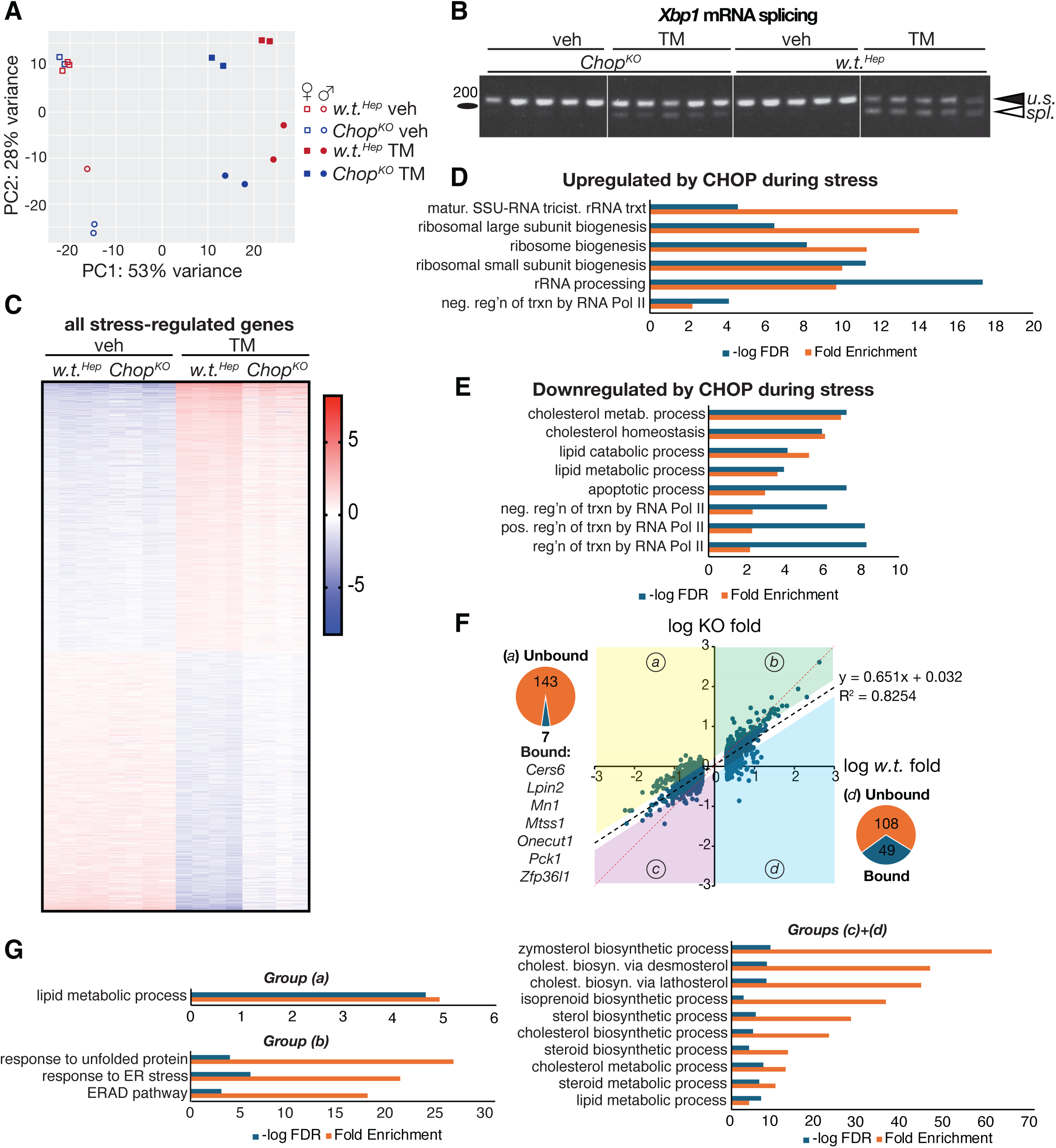
RNA-seq reveals a broad effect of CHOP in augmenting stress-dependent gene regulation **(A)** Principal component analysis of samples used for RNA seq. *Chop^FLuL/FLuL^* animals were exposed to AAV-TBG-GFP or AAV-TBG-FLPo; the former treatment leaves the animals null for CHOP in all tissues (*Chop^KO^*), while the latter restores CHOP competence specifically in hepatocytes (*w.t.^Hep^*). Genotype is indicated by color, sex by shape, and TM treatment by fill. **(B)** *Xbp1* RNA splicing from a larger cohort of animals including the 16 used for RNA-seq **(C)** A heatmap showing all genes significantly regulated (FDR<0.05) more than 2-fold by tunicamycin in *w.t.^Hep^*livers, ordered by magnitude of regulation with upregulated genes on top **(D, E)** Pathway analysis of genes that were lower (D) or higher (E) in their expression in TM-treated *Chop^HKO^* livers compared to wild-type TM-treated. (F) Genes from (C) were plotted for log_10_ induction by TM in *w.t.^Hep^* versus *Chop^KO^* livers. Dashed black line shows the best-fit line and associated equation while the dashed red line shows a hypothetical fit if CHOP deletion had no effect on gene expression. Colored quadrants show genes different in their expression 1.5-fold or greater compared to expected expression from the equation. Pie charts show percentage of genes directly bound by CHOP (from ChIP-seq analysis in Figure 4) or not bound in groups (*a*) and (*d*) with all 7 directly bound genes in group (a) listed below. (G) Pathway analysis of genes from groups (*a*), (*b*), and (*c+d*) in panel F

A TM challenge causes mRNA upregulation and downregulation in the liver in roughly equal measure (Fig. 3C). By taking advantage of previously published microarray analyses after *in vivo* TM challenge in animals lacking PERK, IRE1α, or ATF6α (Teske et al. 2011; Rutkowski et al. 2008; Zhang et al. 2011), we were able to assign the large majority of regulated genes to one or more of the three canonical UPR pathways. Most (∼80%) TM-upregulated genes were regulated by the PERK pathway, over half of which were regulated by PERK alone (Fig. S3A). This trend was even more pronounced among suppressed genes, with almost 90% dependent on PERK and three-fourths of that group dependent on PERK alone (Fig. S3A). Therefore, factors affecting the PERK axis of the UPR stand to have far-reaching effects on stress-dependent gene expression.

A heatmap of all stress-regulated genes showed no appreciable effect of CHOP absent a stress, but a broad diminution of UPR-regulated gene expression—both upregulated genes and downregulated genes—when CHOP was deleted (Fig. 3C). CHOP had the strongest stimulatory effect on pathways of ribosome biogenesis (Fig. 3D), consistent with its previously described role in protein synthesis (Han et al. 2013). Conversely, the pathways most suppressed by CHOP mainly comprised lipid metabolism and Pol II-directed transcription (Fig. 3E). We next plotted the expression of all genes regulated by stress in CHOP-expressing *Chop^fl/fl^* animals versus in CHOP-incompetent *Chop^FLuL/FLuL^* animals. This analysis showed a strong linear correlation (r^2^ ∼ 0.83) with a slope of 0.651 (Fig. 3F), rather than the slope of 1.0 that would be expected were CHOP to have no effect on gene expression. This finding suggests that the most obvious effect of CHOP is a general exacerbation of ER stress, such that cells increase UPR signaling to compensate for its effects.

Because of the apparent general effects of CHOP on ER stress—and with it, the enhancement of transcriptional regulation—we wished to identify genes and pathways upon which CHOP had a larger- or smaller-than-expected effect after this general influence was accounted for. To do this, we identified genes that fell more than 0.176 log_10_ units above or below the regression line from Fig. 3F—i.e., that were 1.5-fold higher (*a, b*) or lower (*c, d*) in their expression than predicted from the regression analysis alone (colored regions Fig. 3F; Worksheet S2).

Genes whose suppression was more strongly dependent on CHOP than expected—group *a*—included those involved in lipid metabolic processes (Fig. 3G). Thus, at the genetic level, CHOP has an outsized effect on suppression of lipid metabolism. Genes for which the converse was true—that CHOP had an outsized effect on their stimulation, i.e., group *d*—did not identify any statistically significant pathway after correcting for false discovery, but groups *c* and *d* together were highly enriched in genes involved in cholesterol metabolism (Fig. 3G). Therefore, deletion of CHOP diminishes the expression of genes in cholesterol metabolic pathways. Group *b*—those genes whose upregulation was not as compromised by CHOP deletion as expected—included a number of classical UPR and ERAD targets (Fig. 3G). In marked contrast to most upregulated genes, the majority of these were regulated by IRE1α or ATF6α or both (Fig. S3B). This genome-level finding is consistent with the observation from targeted qRT-PCR that targets of the IRE1α and ATF6α pathways were relatively indifferent to the presence or absence of CHOP (Fig. 2J, K). Together, these data suggest that CHOP enhances ISR-dependent gene expression with comparatively less effect on the other two branches of the UPR.

### CHOP suppresses hepatic transcription master regulators

CHOP has been proposed to upregulate genes both directly by promoter binding and indirectly by squelching of DNA binding of other transcription factors (Ron and Habener 1992; Ubeda et al. 1996). To identify directly regulated genes, we performed ChIP-seq on pooled livers from three TM-challenged wild-type animals. Approximately 25% of all CHOP-bound sequences were within 3 kb of a transcriptional start site (TSS), substantially exceeding the expected abundance of TSS-flanking sequence in the mammalian genome (Schaefer et al. 2010; Consortium 2012) (Fig. 4A; Worksheet S3). Among all CHOP binding sites, a consensus sequence first identified as a target for CHOP-ATF4 dimers in MEFs was the most highly enriched (Han et al. 2013), followed next by different permutations of binding sites for C/EBP family members including one previously identified for CHOP-C/EBP dimers (Ubeda et al. 1996) (Fig. 4B; Worksheet S4). The 16 highest-ranked (by FDR) motifs belonged to bZIP transcription factors, consistent with the known propensity of these factors to form heterodimers (Miller 2009). This pattern is highly similar to that observed in cultured cells previously (Osman et al. 2023), raising confidence in the reliability of the ChIP-seq results.

**Figure 4:**
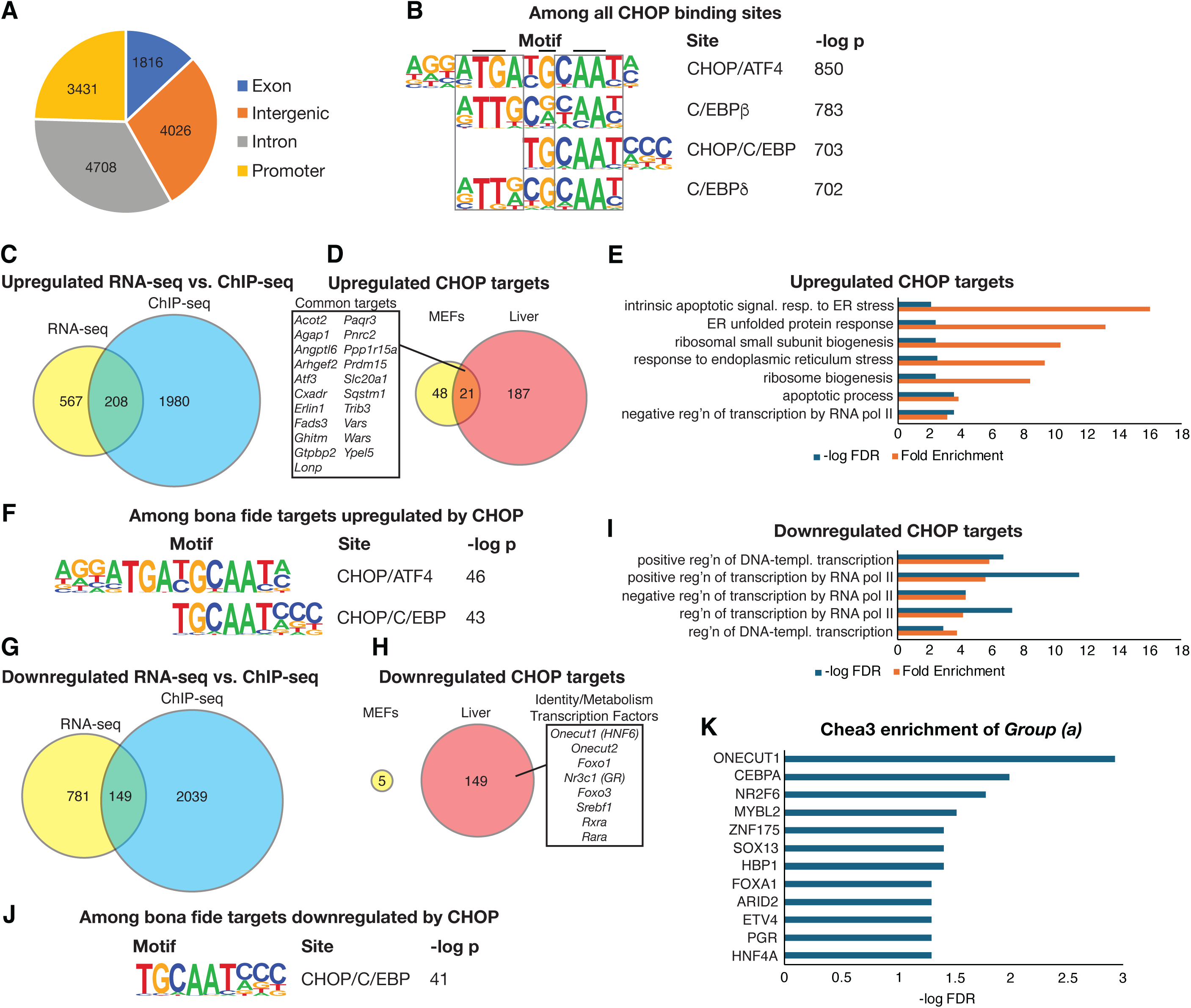
ChIP-seq identifies liver-specific CHOP target genes **(A)** Genome-wide frequency of CHOP binding to the indicated DNA regions, with promoters defined as within 3 kb of a transcriptional start site **(B)** The most enriched known motif binding sites for CHOP genome-wide **(C)** Overlap between promoters bound by CHOP and genes significantly more highly expressed in *w.t.^Hep^* livers after TM challenge than in *Chop^KO^*livers **(D)** Overlap between the 208 *bona fide* CHOP targets for upregulation shown in (C) and similarly identified targets in Han et al.(Han et al. 2013) **(E)** Pathway analysis of *bona fide* targets from (C) **(F)** Motif analysis of *bona fide* targets from (C) **(G-J)** Similar to (C-F) except for CHOP targets for suppression **(K)** ChEA3 analysis to identify enriched transcription factor binding sites among genes in group (*a*) from Figure 3F

We then compared the set of genes bound by CHOP (within 3 kb of the transcriptional start site (TSS) with >10-fold enrichment over input) with those whose expression was significantly different between TM-treated wild-type versus CHOP-incompetent livers. This analysis yielded 208 upregulated and 149 downregulated genes that are likely *bona fide* CHOP targets (Fig. 4C, G). Importantly, a previous study in mouse embryonic fibroblasts (MEFs) identified 69 genes that were likely *bona fide* CHOP targets (Han et al. 2013), and 21 of these were also identified in the liver, including the well-characterized CHOP targets *Ppp1r15a* (*Gadd34*), *Trib3*, *Atf3*, and *Wars* (Fig. 4D). Pathways enriched among *bona fide* upregulated targets included apoptosis (though it is worth reemphasizing that TM at the dose used elicits very little cell death in the liver (Fig. 1C)), ER stress pathways, and ribosome biogenesis (Fig. 4E)—pathways largely consistent with the main functions ascribed to CHOP in MEFs. The most significantly enriched binding sequence for upregulated targets was that of CHOP-ATF4 dimers (Fig. 4F), again consistent with previous results in MEFs (Han et al. 2013).

This comparison supports that the identified genes are likely *bona fide* CHOP targets, and also illustrates that a large fraction of identified targets for upregulation are possibly unique to the liver. By contrast, the previous analysis in MEFs revealed very few targets for direct suppression by CHOP (Han et al. 2013), whereas there were many such putative targets in the liver (Fig. 4G). This finding suggests that direct repression of gene expression by CHOP is contingent on liver-specific gene regulatory pathways in a way that CHOP-dependent upregulation is not. In contrast to upregulated genes, the *bona fide* targets for CHOP-dependent downregulation mostly comprised transcriptional regulators, including several known to regulate hepatocyte differentiation and metabolism (Fig. 4H, I). These included *Onecut1* (*Hnf6*) and *Onecut2*, *Foxo1*, and others. Strikingly, the consensus sequence for CHOP-ATF4 dimers was not identified as enriched among downregulated targets—perhaps unsurprising given that this interaction is thought to stimulate transcription. Rather, the putative CHOP-C/EBP binding site was the most significantly enriched (Fig. 4J).

We considered again the genes whose regulation was more strongly dependent on CHOP than CHOP’s effect on ER stress alone would predict. Unsurprisingly, among genes whose *upregulation* was most strongly enhanced by CHOP (group *d*, Fig. 3F), a substantial number (31%) of those genes’ promoters were directly bound by CHOP, including *Gadd34*. In contrast, only a tiny fraction (<5%) of genes whose *suppression* was strongly enhanced by CHOP (group *a*) were directly bound. Included in these few genes was the transcription factor *Onecut1* (*Hnf6*), which is a known master regulator of hepatocyte differentiation and metabolism (Tian et al. 2025). Interestingly, when the genes from group *a* were analyzed for enriched known transcription factor binding sites, the most significant hit was for ONECUT1 (Fig. 4K). This finding raises the possibility that direct suppression of ONECUT1 by CHOP has a cascading effect on ONECUT1-dependent gene regulation.

### Most novel CHOP direct targets are conserved ex vivo

We next selected CHOP targets of interest for further analysis among those either directly or indirectly regulated by CHOP. Among the former group included three *bona fide* targets from group *d* in Fig. 3E—*Wwtr1*, *Npc1*, and *Tnfrsf12a*—and three targets from group *a*—*Lpin2*, *Onecut1*, and *Pck1*. *Wwtr1* (*Taz*) is a Hippo pathway transcription factor that drives proliferation and dedifferentiation (Ye et al. 2024), *Npc1* regulates lysosomal cholesterol egress and is mutated in Niemann-Pick Disease (Garver et al. 2007), and *Tnfrsf12a* (TWEAK) is an inflammatory cytokine promoting proliferation of hepatocyte progenitors (Jakubowski et al. 2005). *Lpin2* regulates triglyceride synthesis and is mutated in Majeed Syndrome (Zhang and Reue 2017); *Pck1* encodes the committed enzyme of hepatic gluconeogenesis (Yu et al. 2021); and *Onecut1*, as mentioned, transcriptionally regulates hepatocyte differentiation and metabolism (Tian et al. 2025). Validating our RNA-seq results from *Chop^KO^*versus *w.t.^Hep^* livers, each of these genes was, as expected, significantly different in expression between TM-treated *Chop^fl/fl^*to *Chop^HKO^* livers (Fig. 5A). This dependence was also maintained, except for *Tnfrsf12a*, in primary hepatocytes in which CHOP was deleted by Adenoviral delivery of CRE in response to a tunicamycin challenge (Fig. 5B). Largely similar results were obtained when hepatocytes were challenged with the mechanistically distinct ER stressor thapsigargin (TG) (Fig. 5C), suggesting that ER stress more broadly, rather than just TM specifically, elicits this regulation by CHOP.

**Figure 5:**
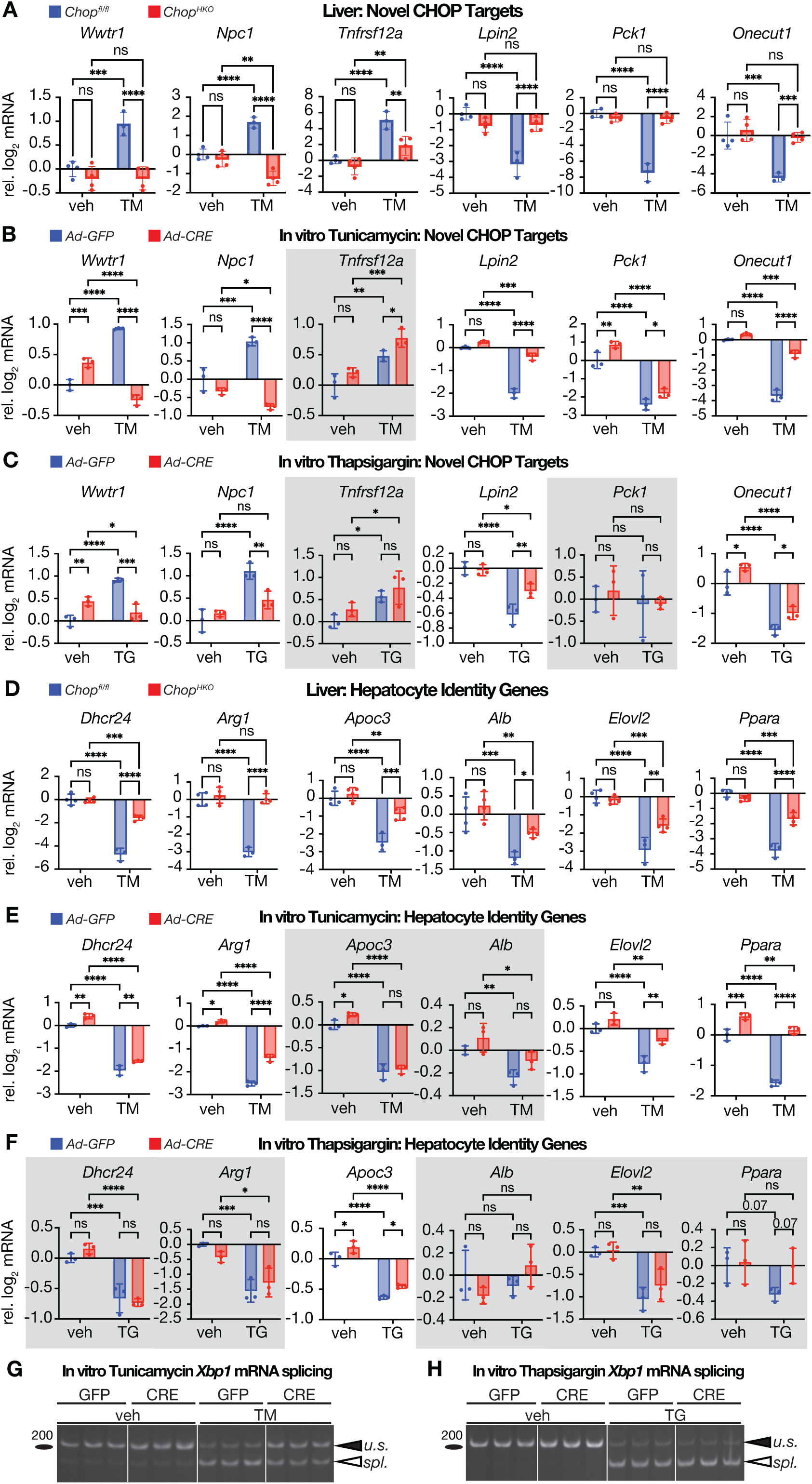
Weakening of CHOP-dependent regulation in primary hepatocytes *in vitro* **(A-C)** Select newly-identified bona fide targets of CHOP were analyzed by qRT-PCR for their expression in the liver after an 8h TM challenge (A), or in *Chop^fl/fl^* primary hepatocytes treated with Ad-CRE to delete *Chop* or Ad-GFP as a control, after an 8h challenge with 5 μg/ml TM (B) or 500 nM TG (C). Gray boxes indicate genes that do not show similar significant CHOP-dependent differences after ER stress challenge *in vitro* as *in vivo*. **(D-F)** Similar to (A-C), except examining genes involved in hepatocyte identity that are not direct targets of CHOP **(G, H)** *Xbp1* mRNA splicing after *in vitro* TM or TG treatment as above

For indirect targets of CHOP, we chose metabolic and hepatocyte identity genes that we had previously characterized as strongly suppressed by ER stress in the liver (Shah et al. 2023)— *Dhcr24*, *Arg1*, *Apoc3*, *Alb*, *Elovl2*, and *Ppara*. Indeed, the suppression of each of these genes was attenuated by CHOP deletion *in vivo* (Fig. 5D). However, *in vitro* their suppression by TM was less robust, with only some genes retaining their CHOP-dependence (Fig. 5E). This result is consistent with the idea that hepatocyte isolation and culture *ex vivo* fundamentally alters the hepatocyte gene regulatory network (Dall et al. 2025), on which CHOP’s effect on these genes is presumably dependent. Supporting this idea, isolated primary hepatocytes had substantially lower expression (often by 10-fold or more) of many genes encoding both master regulators of hepatocyte identity and metabolism, and of downstream hepatocyte identity genes compared to the liver *in vivo* (Fig. S4). Moreover, only one of these indirectly-regulated genes was significantly different in expression between genotypes after TG treatment—and that gene, *Apoc3*, did not differ when measured by fold-change rather than expression (Fig. 5F). This finding was consistent with the observation that CHOP exacerbates ER stress during TM treatment (as seen in *Xbp1* splicing—Fig. 5G) but not during TG treatment (Fig. 5H). This discrepancy is likely attributable to the fact that CHOP exacerbates ER stress by promoting protein synthesis, which TM requires to elicit stress, but which TG does not.

### The effects of CHOP on UPR signaling mirror those seen during chronic stress caused by ATF6α deletion

The above data highlight several impacts of CHOP on ER stress signaling. In the presence of CHOP, IRE1 signaling—presumably a consequence of exacerbated ER stress—is amplified, as seen by *Xbp1* mRNA splicing and RIDD activity (Figs. 2I, 2J, and 3B). Yet the strongest CHOP-dependent gene regulation is seen in ISR-regulated genes, including the metabolic and hepatocyte identity genes that are suppressed by the ISR, in contrast to transcriptional targets of XBP1 and ATF6, which are comparatively unaffected (Figs. 2J, 2K, S3B). This regulation occurs under conditions when eIF2α phosphorylation is diminished (Fig. 2F), but ATF4 expression is not (Fig. S2A). These findings lead to the hypothesis that, by exacerbating ER stress, CHOP elicits a transition in UPR signaling dynamics that depends on ongoing stress, and that prioritizes continued regulation of ISR target genes, including suppression of genes involved in hepatocyte metabolism and identity.

To test this hypothesis, we induced ER stress in animals with hepatocyte-specific deletion of the UPR stress sensor ATF6α. Previous studies in both cultured cells and animals with a whole-body deletion of ATF6α have shown that the ability to effectively restore ER homeostasis after a stress challenge is compromised when ATF6α is deleted (Cao et al. 2013; Wu et al. 2007; Yamamoto et al. 2007). This failure arises because most of the transcriptional targets of ATF6α are ER chaperones and other factors that facilitate ER protein folding and degradation (Yamamoto et al. 2007; Wu et al. 2007). Thus, in its absence, the UPR is unable to fully augment ER protein processing capacity, leading to ongoing, or chronic, stress. If CHOP indeed elicits a transition from acute to chronic UPR signaling, then its effects should be mirrored by loss of ATF6α.

AAV-TBG-CRE was used to delete *Atf6α* specifically in hepatocytes of *Atf6α^fl/fl^* mice (Engin et al. 2013) (*Atf6α^HKO^*), with AAV-TBG-GFP as a control. Animals were then challenged with TM, and livers were collected 48 hours later (Fig. 6A). This longer time point was used because cells and tissues lacking ATF6α initially respond to stress similarly to wild-type, and it is only at later times, when the UPR transitions from non-transcriptional to transcriptional pathways of adaptation, that the failure to fully upregulate ER protein processing factors results in an increasingly profound exacerbation of stress (Rutkowski et al. 2008; Wu et al. 2007). As previously reported in animals with a whole-body deletion of ATF6α, livers in *Atf6α^HKO^* animals were profoundly steatotic after challenge, with wild-type livers only mildly so (Fig. 6B, C). *Xbp1* mRNA splicing was also more persistent in *Atf6α^HKO^* livers (Fig. 6D). Yet eIF2α phosphorylation was much more pronounced in wild-type livers, even though CHOP—an ISR target—was only seen in *Atf6α^HKO^* livers (Fig. 6E). mRNA expression of the ISR targets *Ppp1r15a*/*Gadd34* (∼8-fold), *Trib3* (∼3-4-fold), and *Ero1a* (∼2-fold) was significantly elevated (Fig. 6G). *Chop* mRNA was not significantly different, probably because ATF6α transcriptionally coregulates *Chop* (Ma et al. 2002); the persistence of CHOP protein probably reflects translational stimulation (Palam, Baird, and Wek 2011). The XBP1 transcriptional target *Sec61a1* was modestly elevated (by ∼25%), and *Sec61b* was not significantly different, but the canonical RIDD target *Bloc1s1* was strongly suppressed in *Atf6α^HKO^* livers (Fig. 6H). Furthermore, all of the metabolic genes characterized as indirectly suppressed by CHOP in Fig. 5 were strongly suppressed specifically in *Atf6α^HKO^* livers (Fig. 6I). Lastly, expression of ATF4 was observed in knockout but not wild-type livers (Fig. 6J) despite levels of eIF2α phosphorylation that were elevated in wild-type livers but not elevated in knockouts.

**Figure 6:**
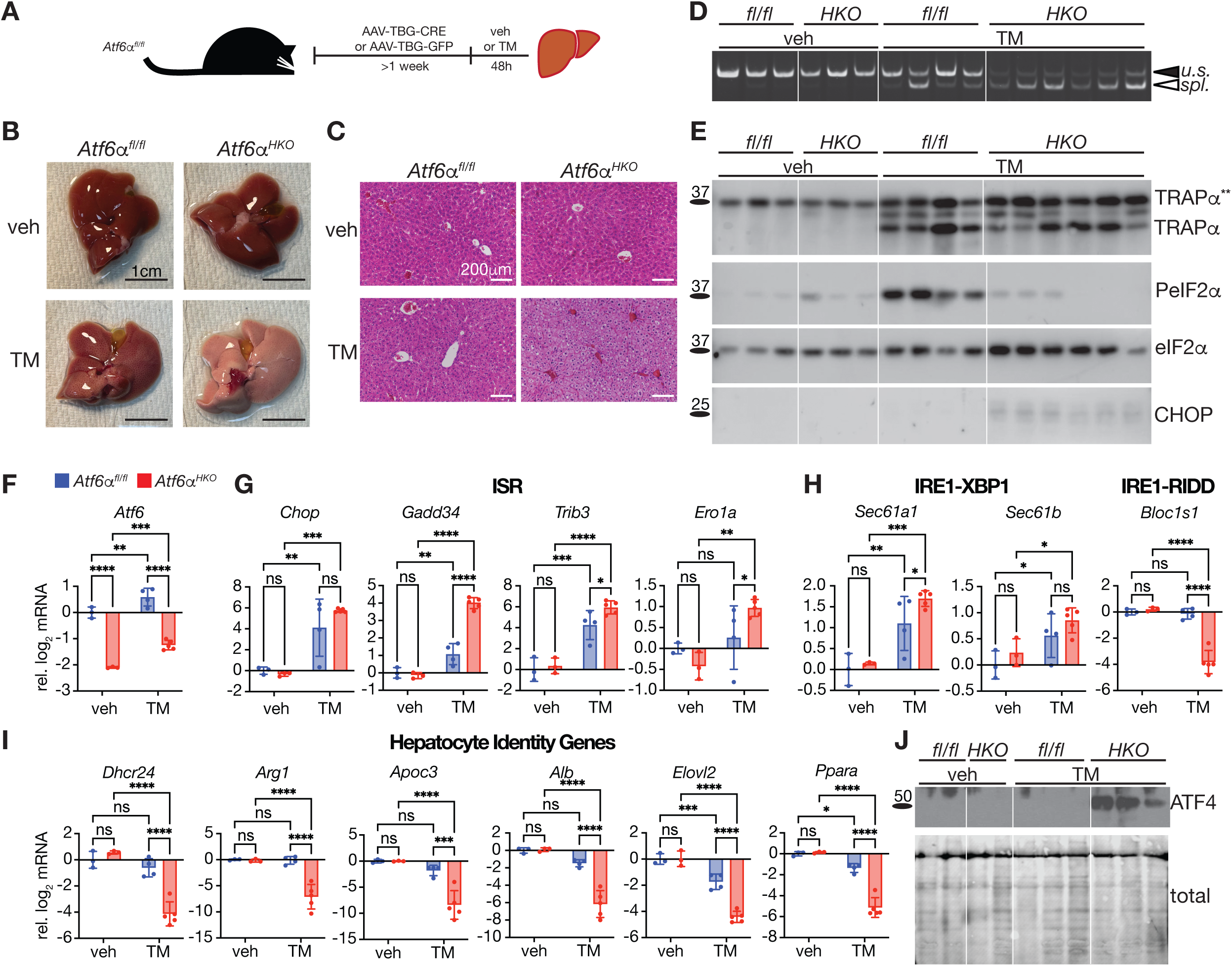
Hepatocyte deletion of ATF6α exacerbates ER stress and permits eIF2α-independent ISR signaling **(A)** Experimental paradigm for *Atf6α* deletion in *Atf6α^fl/fl^* animals and challenge with 1 mg/kg TM for 48h. **(B)** *Xbp1* mRNA splicing **(C)** Whole livers **(D)** Hematoxylin and eosin staining showing extensive microvesicular steatosis in *Atf6α^HKO^* livers after challenge **(E)** Immunoblot to detect CHOP, phosphorylated and total eIF2α, and TRAPα**(F-I)** qRT-PCR quantification of the indicated genes including *Atf6* itself (F), ISR targets (G), IRE1-XBP1 and RIDD targets (H), and hepatocyte identity genes (I) **(J)** Detection of ATF4 in purified nuclei, along with a total protein stain

## Discussion

Our work suggests a previously unappreciated role for CHOP in promoting the transition from first-phase acute to second-phase chronic ER stress signaling *in vivo*. The second phase is characterized by persistent ISR activation despite attenuated eIF2α phosphorylation. A consequence of this transition in the liver is heightened steatosis. We infer this role based on commonalities between the effects that CHOP has on UPR signaling and those that are caused by deletion of ATF6α, which is known to impair the ability of the UPR to efficiently resolve ER stress(Rutkowski et al. 2008; Wu et al. 2007; Yamamoto et al. 2007). Specifically, both the action of CHOP and the loss of ATF6α: (1) lead to prolonged ER stress as suggested by *Xbp1* splicing (Figs. 2I, 3B, and 6D) and also by the global augmentation of stress-dependent gene expression (Figs. 3C, F and (Arensdorf et al. 2013)); (2) promote ATF4 production despite eIF2α dephosphorylation (Figs. S2A and 6J); (3) enhance expression of ISR target genes and activation of RIDD with comparatively less of an effect on expression of XBP1-dependent (and, in *Chop^HKO^*animals, ATF6α-dependent) gene expression (Figs. 2H, J, K, and 6G, H); and (4) magnify the stress-dependent suppression of hepatocyte identity genes (Fig. 5D, 6I).

From our results, we propose the following model (Fig. 7): During the initial exposure to ER stress, phosphorylation of eIF2α by PERK leads to global inhibition of protein synthesis, helping to alleviate the stress burden. At the same time, phosphorylation of eIF2α by PERK translationally stimulates both ATF4 (Harding et al. 2000) and CHOP (Palam, Baird, and Wek 2011), which cooperate transcriptionally to promote eIF2α dephosphorylation and the resumption of protein synthesis (Marciniak et al. 2004; Han et al. 2013). In addition, ATF4 and the other two transcription factors of the UPR—XBP1 and ATF6α—upregulate genes encoding factors that facilitate ER protein folding and cellular health (Lee, Iwakoshi, and Glimcher 2003; Wu et al. 2007), but that would require resumption of protein synthesis for their expression. Further, uniquely in the liver, activation of the ISR also broadly represses the hepatocyte gene regulatory network, leading to suppression of genes involved in hepatocyte identity and metabolism.

**Figure 7:**
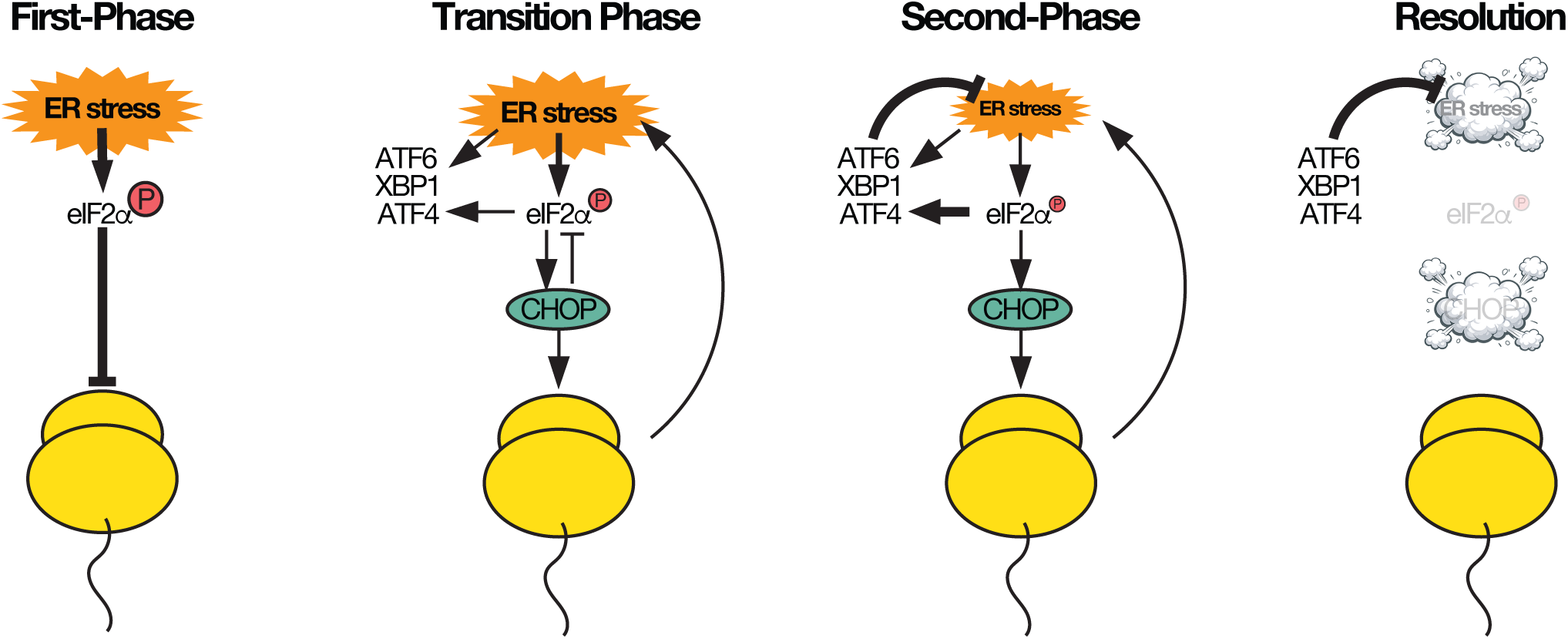
Simplified schematic figure depicting the role of CHOP in mediating the transition from first-phase to second-phase stress signaling. See discussion for details.

The action of CHOP in restoring protein synthesis results in exacerbated ER stress but diminished eIF2α phosphorylation. In this second chronic phase, ISR-dependent gene expression persists even as eIF2α phosphorylation is reversed, although the mechanism by which this persistence occurs is still under investigation. It has been previously shown that deletion of GADD34 paradoxically results in elevated ATF4 synthesis, particularly at later time points (Novoa et al. 2003). Therefore, although the initial phosphorylation of eIF2α promotes ATF4 synthesis, there must exist pathways to sustain ATF4 expression in the face of eIF2α dephosphorylation at later times. A plausible candidate pathway is through eIF3, which has been shown to mediate the translation of mRNAs with upstream open reading frames such as *Atf4* (Mukhopadhyay, Amodeo, and Lee 2023; Herrmannova et al. 2024; Walsh et al. 2025) and to characterize a chronically activated ISR in cells (Guan et al. 2017). Likewise eIF4E has been shown to contribute additional layers of translational regulation during stress (Chen et al. 2010; Chen et al. 2025). In any case, this transition alone and the chronic stress that accompanies it do not appear to necessarily kill cells, as even in animals lacking ATF6α, cell death in hepatocytes in response to an ER stress challenge is modest (Wu et al. 2007). This observation belies the notion that chronic ER stress necessarily results in cell death. Rather, at least when sufficiently mild, such stress perpetuates ISR signaling, one consequence of which in the liver is transcriptional suppression of hepatocyte identity and metabolism genes. For that reason, the most obvious physiological consequence of both CHOP action and ATF6α deletion in the liver is steatosis.

The strongest effects of CHOP on gene expression in the liver are likely attributable to its exacerbation of ER stress and consequent amplification of ISR signaling. However, we have also identified a number of novel direct targets of CHOP. Included among these are genes encoding transcriptional master regulators of hepatocyte differentiation and metabolism including *Onecut1*, *Foxo1*, and others (Fig. 4H), the regulation of which might amplify the impact of CHOP on the expression of downstream target genes. We believe that most upregulated direct targets of CHOP are probably regulated by the previously described CHOP-ATF4 dimers (Han et al. 2013). However, for genes suppressed by CHOP such as *Onecut1*, the mechanism of regulation remains elusive. Both in our experiments and those of others, CHOP overexpression alone has little or no effect on gene regulation, requiring a contemporaneous stress-dependent signal (Han et al. 2013; Chikka et al. 2013). The strong PERK dependence of the UPR at large (Fig. S3) suggests that this signal is likely part of the ISR. For transcriptional upregulation, the production of ATF4 satisfactorily accounts for this extra signal. However, for transcriptional repression, the more likely scenario is that ISR activation results in production of a repressor that then is functionally strengthened by CHOP. An appealing model would be translational regulation of C/EBPα or β, because both of these factors exist in long, activating and short, repressive forms whose preponderance is regulated by translational control (Calkhoven, Muller, and Leutz 2000). They also both—C/EBPα in particular—play important roles in maintaining the hepatocyte gene regulatory network including regulating metabolic pathways (Takiguchi 1998; Ron and Habener 1992). Moreover, CHOP has long been known to associate with both C/EBPs (Ron and Habener 1992), and the preponderance of C/EBP sequences among CHOP’s binding sites support a role for one or both of these proteins. However, hepatocyte deletion of C/EBPα had no effect on stress-dependent gene expression (Fig. S5), and we also found no compelling evidence that ER stress alters the ratio of long to short forms of C/EBPα or β upon ER stress in the liver (data not shown). Therefore, the molecular mechanism by which CHOP represses genes remains to be determined.

Although CHOP clearly promotes cell death during stress, the idea that this is the sole—or even the principal—cell biological consequence of CHOP action merits reevaluation. During stresses modest enough and transient enough to be quickly overcome non-transcriptionally through the rapid inhibition of protein synthesis and the IRE1-dependent decay of ER-associated mRNAs, the eventual resumption of protein synthesis by CHOP presumably has no effect on cell death and rather facilitates the return of cells to the normal levels of biosynthesis needed to sustain function. Yet we propose that CHOP’s actions are of greatest consequence in cells in which ER homeostasis has not been restored by the time that eIF2α dephosphorylation begins. For cells overwhelmingly burdened, it is almost certainly true that CHOP promotes their death. However, during stresses of physiological intensity—for instance, metabolic fluxes (Pfaffenbach et al. 2010)—such cells presumably make up at most a small fraction of the population; the large majority of cells could presumably benefit from an extended window to produce the protective factors that the three limbs of the UPR regulate.

That CHOP would promote both adaptation and death suggest that CHOP is part of a switch mechanism in the UPR, since these two outcomes are mutually exclusive. And, indeed, CHOP exhibits essential features of a switch regulator (Ferrell 1999). Cellular switches must include positive feedback, which CHOP facilitates at two levels: both by promoting the perpetuation of ISR signaling, of which CHOP itself is a part, and by stimulating its own expression in conjunction with ATF4 (Han et al. 2013). A switch also requires a non-linear component, often encoded through a feed-forward mechanism. In the case of the ISR, ATF4, CHOP, and GADD34 are all translationally stimulated by eIF2α phosphorylation, ATF4 transcriptionally regulates *Chop* and *Gadd34*, and CHOP transcriptionally regulates *Gadd34* (Lee, Cevallos, and Jan 2009; Palam, Baird, and Wek 2011; Harding et al. 2000; Ma et al. 2002; Ma and Hendershot 2003; Marciniak et al. 2004). Finally, switches exhibit bistable behavior—i.e., in individual cells, the switch must exist in the “on” or “off” state. We have previously shown both *in vitro* and in the liver *in vivo* that, after CHOP expression peaks uniformly across the cell population, with further time individual cells split into CHOP “on” and CHOP “off” populations (Liu et al. 2024). Based on this behavior, we propose that CHOP is a cell fate switch actuator, and that cell death describes only one outcome of this function. In MEFs, the consequence of the adaptive role of CHOP is cell proliferation (Liu et al. 2024). In the liver, where hepatocytes are not usually proliferating, the consequence is instead the facilitated transition of the ISR/UPR from the acute first-phase to the chronic second-phase.

We propose a beneficial role for CHOP in promoting hepatocyte adaptation to stress despite the fact that the most obvious consequences of CHOP in the liver are steatosis and suppression of hepatocyte identity and metabolism genes. We have previously shown that this genetic suppression is surprisingly beneficial in restoring hepatocyte ER homeostasis, even if detrimental to the liver at large (Shah et al. 2023). This regulation might represent a pathway by which CHOP could contribute to liver injury through a pathway other than cell death, even in contexts where CHOP-mediated cell death occurs. Physiological *in vivo* challenges in which the fate of individual cells can be followed will allow the cell biological contributions of CHOP (death versus loss of identity versus return to homeostasis) to be dissected.

## Materials and Methods

### Animals

All animal experiments were approved by and in accordance with Institutional Animal Care and Use Committee procedures. Mice were kept on a 12h light/dark cycle in the University of Iowa Animal Care Facility. Creation of *Chop^FLuL/FLuL^*mice and breeding of *Chop^fl/fl^* mice is described in (Liu et al. 2024). *Cebpa^fl/fl^* and *Atf6α^fl/fl^* mice were obtained from Jackson labs (strains 006447 and 028253 respectively) and are described (Zhang et al. 2004; Engin et al. 2013). Animals of both sexes, 8-14 weeks of age, were used. Viral deletions were performed with 1-2 x 10^11^ vg/animal of AAV-TBG-CRE, AAV-TBG-FLPo, or AAV-TBG-EGFP, obtained from either Vector Biolabs or the University of Iowa Viral Vector Core. Virus was injected intraperitoneally and experiments were performed at least 1 week after injection to allow for complete deletion. TM from stock was diluted in PBS and injected intraperitoneally at 1 mg/kg b.w. Similarly-diluted DMSO was used as vehicle. Animals were fasted 4h prior to euthanasia, which was by cervical dislocation. Efficacy of TM treatment was confirmed by immunoblot for the glycoprotein TRAPα, and efficacy of genetic deletions was confirmed by qRT-PCR, immunoblot, or both. Any animal for which either treatment was inefficacious was excluded from further analysis. For acetaminophen (APAP) challenge, APAP was dissolved to 30 mg/ml in warm PBS prior to injection at 300 mg/kg. Primary hepatocytes were isolated and cultured as described (Gansemer et al. 2020). After isolation, hepatocytes were allowed to rest 4h, then exposed overnight to Ad-CRE or Ad-GFP at an M.O.I. of 5. Treatments with ER stressors were then as described in the text.

### Molecular and Cellular Analyses

qRT-PCR was carried out as described (Rutkowski et al. 2006). Briefly, all primer pairs were first validated for specificity and quantitative efficiency using standard curves, melt curves, and no-RT controls. iScript (Bio-Rad) was used for cDNA synthesis, using 0.4 μg of Trizol-purified RNA in 8 μl reactions, followed by 1/100 dilution and PCR using iTaq SYBR Green (Bio-Rad). Reaction conditions were 40 cycles of 94° 10” and 60° 45”. qRT-PCR primer sequences are as follows:

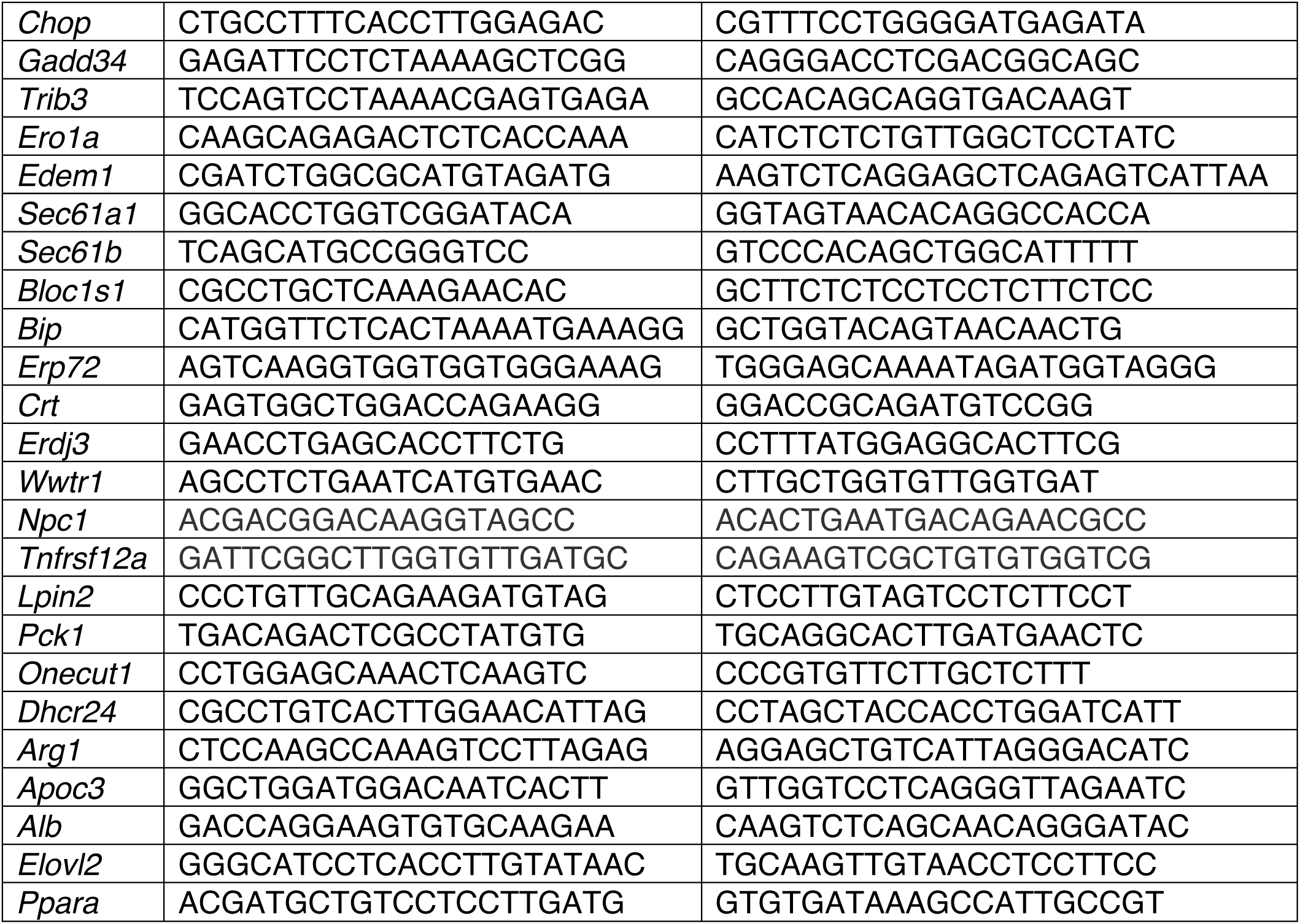
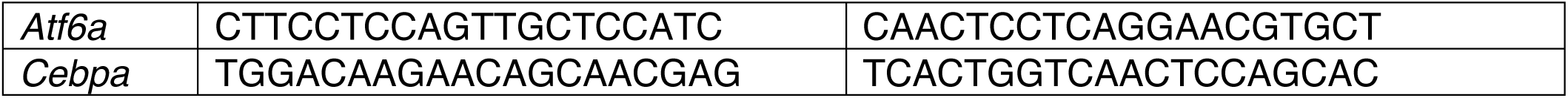

For *Xbp1* RT-PCR, Superscript III One-Step RT-PCR system (Thermo) was used with primers TTGTGGTTGAGAACCAGG and TCCATGGGAAGATGTTCTGG, and products were separated on a 10% acrylamide gel. Immunoblots were carried out using overnight tank transfer onto PVDF membrane and using the ECL Prime Detection kit (Cytiva). Membranes were never stripped and reprobed—rather, comigrating samples were detected using duplicate samples run in parallel. Antibodies were as follows:

|  |  |  |
| --- | --- | --- |
| CNX | ADA-SPA-865 | Assay Designs |
| CHOP | 15204-1-AP | Proteintech |
| TRAPα | N/A | Gift from R.S. Hegde |
| BiP | 610978 | BD Transduction |
| P-eIF2α | 3398 | Cell Signaling Technology |
| Tot-eIF2α | 9722 | Cell Signaling Technology |
| ATF4 (Ab1) | sc200 (discontinued) | Santa Cruz |
| ATF4 (Ab2) | 11815 | Cell Signaling Technology |
| H2A | 2578 | Cell Signaling Technology |

Antibodies against CHOP and ATF4 were validated for specificity using knockout cell lines and/or animals. Each ATF4 antibody was found to also produce numerous non-specific bands in whole liver lysates (different between the two antibodies), including bands of similar migration to the correct species identified by both antibodies in nuclei. P-eIF2α antibody was validated based on comigration with tot-eIF2α and robust increase upon ER stress challenges. CNX antibody was validated by molecular weight of full-length protein and of the membrane-protected ER lumenal fragment after protease digestion of ER-derived microsomes. TRAPα antibody was validated by molecular weight, correct glycan shifting upon TM treatment, and ER-localized immunofluorescence. BiP antibody was validated by protein overexpression, ER-localized immunofluorescence, and by protected ER lumenal disposition upon protease digestion in rough microsomes. Tot-eIF2α antibody was validated by comigration with P-eIF2α band. H2A antibody was validated by molecular weight and its presence in nuclear extracts. Unprocessed blot images are shown in Fig. S6. No-Stain (Invitrogen) was used to detect total protein on membranes.

Triglycerides were measured from liver homogenates using the Colorimetric Triglyceride Assay Kit (Novus) following manufacturer’s instructions. For H&E staining and immunohistochemistry, samples were fixed in 10% formalin, paraffin-embedded, and sectioned. IHC was performed using the Vectastain Elite ABC-HRP kit with ImmPACT DAB Substrate (Vector Laboratories) following the manufacturer’s protocols. Histological and histochemical imaging used a Nikon Eclipse E600 microscope, with slides mounted using a 1:1 mix of Permount and Xylene (Fisher). For fluorescent imaging, liver pieces were fixed overnight in freshly made 4% paraformaldehyde, cryoprotected in 15% and 30% sucrose in PBS 16 and 24 - 48h respectively, and frozen in OCT over dry ice. TUNEL staining used the CF640R TUNEL Assay Kit (Biotium) following the manufacturer’s instructions. Fluorescent imaging used a Zeiss 700 Confocal Microscope. Slides were mounted with Prolong Diamond AntiFade (ThermoFisher). For nuclear purification, frozen liver pieces (∼50 mg) were plunged into ice-cold Buffer A (0.25 M sucrose, 50 mM Hepes, 22mM NaOH, 25 mM KAc, 5 mM MgAc, 1 mM PMSF), removed and quickly minced with a scalpel, returned to 1.25 ml of Buffer A, homogenized with a loose glass-on-glass homogenizer, centrifuged at 4° 800g 10’, the pellet resuspended in 1.25 ml Buffer A, rehomogenized with a tight glass-on-glass homogenizer, and centrifuged again at 4° 800g 10’. The pellet was crudely resuspended in 250 μl Buffer A, to which 500 μl of Buffer B was added (same as Buffer A but with 2.3 M sucrose). Samples were homogenized again and transferred to TLA 120.2 tubes, then underlaid with 250ul of Buffer B. Samples were spun 55,000 rpm, 4°, 60’. Pellets were resuspended in 125 μl Buffer A, homogenized in a small glass-on-glass homogenizer, and flash frozen.

### Deep sequencing

For RNA-seq, Trizol-purified RNAs were re-purified using RNeasy (QIAgen) and RNA integrity validated using NanoDrop. Briefly, 2 μg/sample of RNA was used for sequencing using Illumina HiSeq 4000/75PE Sequencer. 23 million reads were generated for each sample. To quantify transcript-level abundances, we performed the quality control for RNA-seq reads and used the Salmon tool with mapping-based mode again the mouse reference transcriptome (mm10). After transcript quantification, the differential gene expression analysis between different groups was conducted with the DESeq2 package. Differential expression was defined as log_2_fc > 0.58 and FDR-adjusted p-value < 0.05.

ChIP-seq was performed by Zymo Research using pooled samples from 3 mice treated with 1 mg/kg b.w. TM for 8h. IP was with CHOP antibody B-3 (sc7351, Santa Cruz), previously validated for specificity (Chikka et al. 2013), and validated prior to ChIP-seq against control Ig (12-371, Millipore Sigma) for the known CHOP-binding region in the *Gadd34* promoter (Marciniak et al. 2004). Approximately 100-250 μg of chromatin was used. ChIP-seq libraries were prepared using a proprietary library preparation protocol from Zymo Research. ChIP-Seq libraries were diluted to 2 nM before being multiplexed and run on the Next-Gen Sequencing platform –HiSeq using single-end reads. For sequencing data analysis, FastQC was applied to conduct quality control for ChIP-seq data. Reads were aligned to the mouse reference genome mm10 by Bowtie with at most 2 mismatches. Reads that appeared more than twice at the same position on the same strand were discarded to remove PCR duplication. After filtering bam files by Samtools v1.9 and Sambamba v0.7.1, we used MACS2 v2.1.2 to call peaks with default parameters. The R package ChIPseeker was applied to annotate peaks. ChIP-seq motif analysis was performed with HOMER, searching against IMAGE known motifs. Pathway analysis for RNA-seq and ChIP-seq was performed using DAVID (Dennis et al. 2003), and enriched transcription factor binding sites were analyzed using ChEA3 (Keenan et al. 2019) and the ReMap database.

### Statistics

Statistical analysis of directly measured molecular data was by two-way ANOVA comparing against row and column using Prism (GraphPad) with Fisher’s LSD post-hoc analysis and confirmation of Gaussian residuals. For pathway analyses, false discovery rate (FDR) was applied.

### Author contributions

TLV, KL, ZZ, and RCA performed experiments. CZ and HC contributed bioinformatic analysis. KL and DTR conceived the project and designed the experiments. DTR wrote the manuscript. All authors approved the manuscript. AI was used to create the disappearing cloud image in Fig. 7.

## Supporting information

Worksheet S1

Worksheet S2

Worksheet S3

Worksheet S4

## Acknowledgments

This work was funded by NIH grant GM115424 to DTR and by funds from the University of Iowa Department of Anatomy and Cell Biology. Funders had no other role in the project. From the University of Iowa Department of Anatomy and Cell Biology, we thank members of the Ling Yang lab, especially Juan Rodriguez, for technical assistance with primary hepatocyte preparation, and members of the Martine Dunnwald lab, especially Lindsey Rhea and Emily Adelizzi for assistance with histology and imaging. We also thank Eric Van Otterloo for sharing reagents. We thank Udayan Apte (Kansas University Medical Center) for advice on APAP injections and Ramanujan Hegde (MRC Cambridge, UK) for the TRAPα antibody.

## References

1. Adamson, B., T. M. Norman, M. Jost, M. Y. Cho, J. K. Nunez, Y. Chen, J. E. Villalta, L. A. Gilbert, M. A. Horlbeck, M. Y. Hein, R. A. Pak, A. N. Gray, C. A. Gross, A. Dixit, O. Parnas, A. Regev, and J. S. Weissman. 2016. ’A Multiplexed Single-Cell CRISPR Screening Platform Enables Systematic Dissection of the Unfolded Protein Response’, Cell, 167: 1867–82 e21.

2. Arensdorf, A. M., D. Dezwaan McCabe, R. J. Kaufman, and D. T. Rutkowski. 2013. ’Temporal clustering of gene expression links the metabolic transcription factor HNF4alpha to the ER stress-dependent gene regulatory network’, Front Genet, 4: 188.

3. Barone, M. V., A. Crozat, A. Tabaee, L. Philipson, and D. Ron. 1994. ’CHOP (GADD153) and its oncogenic variant, TLS-CHOP, have opposing effects on the induction of G1/S arrest’, Genes Dev, 8: 453–64.

4. Bright, M. D., D. N. Itzhak, C. P. Wardell, G. J. Morgan, and F. E. Davies. 2015. ’Cleavage of BLOC1S1 mRNA by IRE1 Is Sequence Specific, Temporally Separate from XBP1 Splicing, and Dispensable for Cell Viability under Acute Endoplasmic Reticulum Stress’, Mol Cell Biol, 35: 2186–202.

5. Calkhoven, C. F., C. Muller, and A. Leutz. 2000. ’Translational control of C/EBPalpha and C/EBPbeta isoform expression’, Genes Dev, 14: 1920–32.

6. Canales, A., M. Rosinger, J. Sastre, I. C. Felli, J. Jimenez-Barbero, G. Gimenez-Gallego, and C. Fernandez-Tornero. 2017. ’Hidden alpha-helical propensity segments within disordered regions of the transcriptional activator CHOP’, PLoS One, 12: e0189171.

7. Cao, S. S., E. M. Zimmermann, B. M. Chuang, B. Song, A. Nwokoye, J. E. Wilkinson, K. A. Eaton, and R. J. Kaufman. 2013. ’The Unfolded Protein Response and Chemical Chaperones Reduce Protein Misfolding and Colitis in Mice’, Gastroenterology.

8. Chen, C. W., D. Papadopoli, K. J. Szkop, B. J. Guan, M. Alzahrani, J. Wu, R. Jobava, M. M. Asraf, D. Krokowski, A. Vourekas, W. C. Merrick, A. A. Komar, A. E. Koromilas, M. Gorospe, M. J. Payea, F. Wang, B. L. L. Clayton, P. J. Tesar, A. Schaffer, A. Miron, I. Bederman, E. Jankowsky, C. Vogel, L. S. Valasek, J. D. Dinman, Y. Zhang, B. Tirosh, O. Larsson, I. Topisirovic, and M. Hatzoglou. 2025. ’Plasticity of the mammalian integrated stress response’, Nature.

9. Chen, Y. J., B. C. Tan, Y. Y. Cheng, J. S. Chen, and S. C. Lee. 2010. ’Differential regulation of CHOP translation by phosphorylated eIF4E under stress conditions’, Nucleic Acids Res, 38: 764–77.

10. Chikka, M. R., D. D. McCabe, H. M. Tyra, and D. T. Rutkowski. 2013. ’C/EBP homologous protein (CHOP) contributes to suppression of metabolic genes during endoplasmic reticulum stress in the liver’, J Biol Chem, 288: 4405–15.

11. Consortium, Encode Project. 2012. ’An integrated encyclopedia of DNA elements in the human genome’, Nature, 489: 57–74.

12. Costa-Mattioli, M., and P. Walter. 2020. ’The integrated stress response: From mechanism to disease’, Science, 368.

13. Dall, M., B. Stocks, D. T. Cervone, A. S. Deshmukh, and J. T. Treebak. 2025. ’Hepatocyte dedifferentiation in 2D culture reveals extensive transcriptomic and proteomic rewiring’, Hepatol Commun, 9.

14. Dennis, G., Jr., B. T. Sherman, D. A. Hosack, J. Yang, W. Gao, H. C. Lane, and R. A. Lempicki. 2003. ’DAVID: Database for Annotation, Visualization, and Integrated Discovery’, Genome Biol, 4: P3.

15. DeZwaan-McCabe, D., J. D. Riordan, A. M. Arensdorf, M. S. Icardi, A. J. Dupuy, and D. T. Rutkowski. 2013. ’The stress-regulated transcription factor CHOP promotes hepatic inflammatory gene expression, fibrosis, and oncogenesis’, PLoS Genet, 9: e1003937.

16. DeZwaan-McCabe, D., R. D. Sheldon, M. C. Gorecki, D. F. Guo, E. R. Gansemer, R. J. Kaufman, K. Rahmouni, M. P. Gillum, E. B. Taylor, L. M. Teesch, and D. T. Rutkowski. 2017. ’ER Stress Inhibits Liver Fatty Acid Oxidation while Unmitigated Stress Leads to Anorexia-Induced Lipolysis and Both Liver and Kidney Steatosis’, Cell Rep, 19: 1794–806.

17. Engin, F., A. Yermalovich, T. Nguyen, S. Hummasti, W. Fu, D. L. Eizirik, D. Mathis, and G. S. Hotamisligil. 2013. ’Restoration of the unfolded protein response in pancreatic beta cells protects mice against type 1 diabetes’, Sci Transl Med, 5: 211ra156.

18. Ferrell, J. E., Jr. 1999. ’Building a cellular switch: more lessons from a good egg’, Bioessays, 21: 866–70.

19. Galehdar, Z., P. Swan, B. Fuerth, S. M. Callaghan, D. S. Park, and S. P. Cregan. 2010. ’Neuronal apoptosis induced by endoplasmic reticulum stress is regulated by ATF4-CHOP-mediated induction of the Bcl-2 homology 3-only member PUMA’, J Neurosci, 30: 16938–48.

20. Gansemer, E. R., K. S. McCommis, M. Martino, A. Q. King-McAlpin, M. J. Potthoff, B. N. Finck, E. B. Taylor, and D. T. Rutkowski. 2020. ’NADPH and Glutathione Redox Link TCA Cycle Activity to Endoplasmic Reticulum Homeostasis’, iScience, 23: 101116.

21. Garver, W. S., D. Jelinek, J. N. Oyarzo, J. Flynn, M. Zuckerman, K. Krishnan, B. H. Chung, and R. A. Heidenreich. 2007. ’Characterization of liver disease and lipid metabolism in the Niemann-Pick C1 mouse’, J Cell Biochem, 101: 498–516.

22. Guan, B. J., V. van Hoef, R. Jobava, O. Elroy-Stein, L. S. Valasek, M. Cargnello, X. H. Gao, D. Krokowski, W. C. Merrick, S. R. Kimball, A. A. Komar, A. E. Koromilas, A. Wynshaw-Boris, I. Topisirovic, O. Larsson, and M. Hatzoglou. 2017. ’A Unique ISR Program Determines Cellular Responses to Chronic Stress’, Mol Cell, 68: 885–900 e6.

23. Han, J., S. H. Back, J. Hur, Y. H. Lin, R. Gildersleeve, J. Shan, C. L. Yuan, D. Krokowski, S. Wang, M. Hatzoglou, M. S. Kilberg, M. A. Sartor, and R. J. Kaufman. 2013. ’ER-stress-induced transcriptional regulation increases protein synthesis leading to cell death’, Nat Cell Biol, 15: 481–90.

24. Harding, H. P., I. Novoa, Y. Zhang, H. Zeng, R. Wek, M. Schapira, and D. Ron. 2000. ’Regulated translation initiation controls stress-induced gene expression in mammalian cells’, Mol Cell, 6: 1099–108.

25. Harding, H. P., Y. Zhang, H. Zeng, I. Novoa, P. D. Lu, M. Calfon, N. Sadri, C. Yun, B. Popko, R. Paules, D. F. Stojdl, J. C. Bell, T. Hettmann, J. M. Leiden, and D. Ron. 2003. ’An integrated stress response regulates amino acid metabolism and resistance to oxidative stress’, Mol Cell, 11: 619–33.

26. Herrmannova, A., J. Jelinek, K. Pospisilova, F. Kerenyi, T. Vomastek, K. Watt, J. Brabek, M. P. Mohammad, S. Wagner, I. Topisirovic, and L. S. Valasek. 2024. ’Perturbations in eIF3 subunit stoichiometry alter expression of ribosomal proteins and key components of the MAPK signaling pathways’, Elife, 13.

27. Hetz, C., K. Zhang, and R. J. Kaufman. 2020. ’Mechanisms, regulation and functions of the unfolded protein response’, Nat Rev Mol Cell Biol, 21: 421–38.

28. Hollien, J., J. H. Lin, H. Li, N. Stevens, P. Walter, and J. S. Weissman. 2009. ’Regulated Ire1-dependent decay of messenger RNAs in mammalian cells’, J Cell Biol, 186: 323–31.

29. Jakubowski, A., C. Ambrose, M. Parr, J. M. Lincecum, M. Z. Wang, T. S. Zheng, B. Browning, J. S. Michaelson, M. Baetscher, B. Wang, D. M. Bissell, and L. C. Burkly. 2005. ’TWEAK induces liver progenitor cell proliferation’, J Clin Invest, 115: 2330–40.

30. Ji, C., R. Mehrian-Shai, C. Chan, Y. H. Hsu, and N. Kaplowitz. 2005. ’Role of CHOP in hepatic apoptosis in the murine model of intragastric ethanol feeding’, Alcohol Clin Exp Res, 29: 1496–503.

31. Keenan, A. B., D. Torre, A. Lachmann, A. K. Leong, M. L. Wojciechowicz, V. Utti, K. M. Jagodnik, E. Kropiwnicki, Z. Wang, and A. Ma’ayan. 2019. ’ChEA3: transcription factor enrichment analysis by orthogonal omics integration’, Nucleic Acids Res, 47: W212–W24.

32. Kiourtis, C., A. Wilczynska, C. Nixon, W. Clark, S. May, and T. G. Bird. 2021. ’Specificity and off-target effects of AAV8-TBG viral vectors for the manipulation of hepatocellular gene expression in mice’, Biol Open, 10.

33. Krokowski, D., J. Han, M. Saikia, M. Majumder, C. L. Yuan, B. J. Guan, E. Bevilacqua, O. Bussolati, S. Broer, P. Arvan, M. Tchorzewski, M. D. Snider, M. Puchowicz, C. M. Croniger, S. R. Kimball, T. Pan, A. E. Koromilas, R. J. Kaufman, and M. Hatzoglou. 2013. ’A self-defeating anabolic program leads to beta-cell apoptosis in endoplasmic reticulum stress-induced diabetes via regulation of amino acid flux’, J Biol Chem, 288: 17202–13.

34. Lee, A. H., N. N. Iwakoshi, and L. H. Glimcher. 2003. ’XBP-1 regulates a subset of endoplasmic reticulum resident chaperone genes in the unfolded protein response’, Mol Cell Biol, 23: 7448–59.

35. Lee, Y. Y., R. C. Cevallos, and E. Jan. 2009. ’An upstream open reading frame regulates translation of GADD34 during cellular stresses that induce eIF2alpha phosphorylation’, J Biol Chem, 284: 6661–73.

36. Li, G., M. Mongillo, K. T. Chin, H. Harding, D. Ron, A. R. Marks, and I. Tabas. 2009. ’Role of ERO1-alpha-mediated stimulation of inositol 1,4,5-triphosphate receptor activity in endoplasmic reticulum stress-induced apoptosis’, J Cell Biol, 186: 783–92.

37. Liu, K., C. Zhao, R. C. Adajar, D. DeZwaan-McCabe, and D. T. Rutkowski. 2024. ’A beneficial adaptive role for CHOP in driving cell fate selection during ER stress’, EMBO Rep, 25: 228–53.

38. Ma, Y., J. W. Brewer, J. A. Diehl, and L. M. Hendershot. 2002. ’Two distinct stress signaling pathways converge upon the CHOP promoter during the mammalian unfolded protein response’, J Mol Biol, 318: 1351–65.

39. Ma, Y., and L. M. Hendershot. 2003. ’Delineation of a negative feedback regulatory loop that controls protein translation during endoplasmic reticulum stress’, J Biol Chem, 278: 34864–73.

40. Marciniak, S. J., C. Y. Yun, S. Oyadomari, I. Novoa, Y. Zhang, R. Jungreis, K. Nagata, H. P. Harding, and D. Ron. 2004. ’CHOP induces death by promoting protein synthesis and oxidation in the stressed endoplasmic reticulum’, Genes Dev, 18: 3066–77.

41. Miller, M. 2009. ’The importance of being flexible: the case of basic region leucine zipper transcriptional regulators’, Curr Protein Pept Sci, 10: 244–69.

42. Mukhopadhyay, S., M. E. Amodeo, and A. S. Y. Lee. 2023. ’eIF3d controls the persistent integrated stress response’, Mol Cell, 83: 3303–13 e6.

43. Novoa, I., Y. Zhang, H. Zeng, R. Jungreis, H. P. Harding, and D. Ron. 2003. ’Stress-induced gene expression requires programmed recovery from translational repression’, EMBO J, 22: 1180–7.

44. Osman, A., M. Linden, T. Osterlund, C. Vannas, L. Andersson, M. Escobar, A. Stahlberg, and P. Aman. 2023. ’Identification of genomic binding sites and direct target genes for the transcription factor DDIT3/CHOP’, Exp Cell Res, 422: 113418.

45. Palam, L. R., T. D. Baird, and R. C. Wek. 2011. ’Phosphorylation of eIF2 facilitates ribosomal bypass of an inhibitory upstream ORF to enhance CHOP translation’, J Biol Chem, 286: 10939–49.

46. Pennuto, M., E. Tinelli, M. Malaguti, U. Del Carro, M. D’Antonio, D. Ron, A. Quattrini, M. L. Feltri, and L. Wrabetz. 2008. ’Ablation of the UPR-mediator CHOP restores motor function and reduces demyelination in Charcot-Marie-Tooth 1B mice’, Neuron, 57: 393–405.

47. Pfaffenbach, K. T., A. M. Nivala, L. Reese, F. Ellis, D. Wang, Y. Wei, and M. J. Pagliassotti. 2010. ’Rapamycin Inhibits Postprandial-Mediated X-Box-Binding Protein-1 Splicing in Rat Liver’, J Nutr, 140: 879–84.

48. Prigge, J. R., J. A. Wiley, E. A. Talago, E. M. Young, L. L. Johns, J. A. Kundert, K. M. Sonsteng, W. P. Halford, M. R. Capecchi, and E. E. Schmidt. 2013. ’Nuclear double-fluorescent reporter for in vivo and ex vivo analyses of biological transitions in mouse nuclei’, Mamm Genome.

49. Pu, W., H. Zhang, X. Huang, X. Tian, L. He, Y. Wang, L. Zhang, Q. Liu, Y. Li, Y. Li, H. Zhao, K. Liu, J. Lu, Y. Zhou, P. Huang, Y. Nie, Y. Yan, L. Hui, K. O. Lui, and B. Zhou. 2016. ’Mfsd2a+ hepatocytes repopulate the liver during injury and regeneration’, Nat Commun, 7: 13369.

50. Puthalakath, H., L.A. O’Reilly, P. Gunn, L. Lee, P.N. Kelly, N.D. Huntington, P.D. Hughes, E.M. Michalak, J. McKimm-Breschkin, N. Motoyama, T. Gotoh, S. Akira, P. Bouillet, and A. Strasser. 2007. ’ER stress triggers apoptosis by activating BH3-only protein Bim’, Cell, 129: 1337–49.

51. Ron, D., and J. F. Habener. 1992. ’CHOP, a novel developmentally regulated nuclear protein that dimerizes with transcription factors C/EBP and LAP and functions as a dominant-negative inhibitor of gene transcription’, Genes Dev, 6: 439–53.

52. Rutkowski, D. T., S. M. Arnold, C. N. Miller, J. Wu, J. Li, K. M. Gunnison, K. Mori, A. A. Sadighi Akha, D. Raden, and R. J. Kaufman. 2006. ’Adaptation to ER stress is mediated by differential stabilities of pro-survival and pro-apoptotic mRNAs and proteins’, PLoS Biol, 4: e374.

53. Rutkowski, D. T., J. Wu, S. H. Back, M. U. Callaghan, S. P. Ferris, J. Iqbal, R. Clark, H. Miao, J. R. Hassler, J. Fornek, M. G. Katze, M. M. Hussain, B. Song, J. Swathirajan, J. Wang, G. D. Yau, and R. J. Kaufman. 2008. ’UPR pathways combine to prevent hepatic steatosis caused by ER stress-mediated suppression of transcriptional master regulators’, Dev Cell, 15: 829–40.

54. Scaiewicz, V., A. Nahmias, R. T. Chung, T. Mueller, B. Tirosh, and O. Shibolet. 2013. ’CCAAT/enhancer-binding protein homologous (CHOP) protein promotes carcinogenesis in the DEN-induced hepatocellular carcinoma model’, PLoS One, 8: e81065.

55. Schaefer, U., R. Kodzius, C. Kai, J. Kawai, P. Carninci, Y. Hayashizaki, and V. B. Bajic. 2010. ’High sensitivity TSS prediction: estimates of locations where TSS cannot occur’, PLoS One, 5: e13934.

56. Shah, A., I. Huck, K. Duncan, E. R. Gansemer, K. Liu, R. C. Adajar, U. Apte, M. A. Stamnes, and D. T. Rutkowski. 2023. ’Interference with the HNF4-dependent gene regulatory network diminishes endoplasmic reticulum stress in hepatocytes’, Hepatol Commun, 7.

57. Sidrauski, C., D. Acosta-Alvear, A. Khoutorsky, P. Vedantham, B. R. Hearn, H. Li, K. Gamache, C. M. Gallagher, K. K. Ang, C. Wilson, V. Okreglak, A. Ashkenazi, B. Hann, K. Nader, M. R. Arkin, A. R. Renslo, N. Sonenberg, and P. Walter. 2013. ’Pharmacological brake-release of mRNA translation enhances cognitive memory’, Elife, 2: e00498.

58. Singh, V. K., I. Pacheco, V. N. Uversky, S. P. Smith, R. J. MacLeod, and Z. Jia. 2008. ’Intrinsically disordered human C/EBP homologous protein regulates biological activity of colon cancer cells during calcium stress’, J Mol Biol, 380: 313–26.

59. Sood, R., A. C. Porter, K. Ma, L. A. Quilliam, and R. C. Wek. 2000. ’Pancreatic eukaryotic initiation factor-2alpha kinase (PEK) homologues in humans, Drosophila melanogaster and Caenorhabditis elegans that mediate translational control in response to endoplasmic reticulum stress’, Biochem J, 346 Pt 2: 281–93.

60. Takiguchi, M. 1998. ’The C/EBP family of transcription factors in the liver and other organs’, Int J Exp Pathol, 79: 369–91.

61. Teske, B. F., S. A. Wek, P. Bunpo, J. K. Cundiff, J. N. McClintick, T. G. Anthony, and R. C. Wek. 2011. ’The eIF2 kinase PERK and the integrated stress response facilitate activation of ATF6 during endoplasmic reticulum stress’, Mol Biol Cell, 22: 4390–405.

62. Tian, M., W. Gao, S. Ma, H. Cao, Y. Zhang, F. An, J. Qi, and Z. Yang. 2025. ’Role of HNF6 in liver homeostasis and pathophysiology’, Mol Med, 31: 48.

63. Ubeda, M., X. Z. Wang, H. Zinszner, I. Wu, J. F. Habener, and D. Ron. 1996. ’Stress-induced binding of the transcriptional factor CHOP to a novel DNA control element’, Mol Cell Biol, 16: 1479–89.

64. Uzi, D., L. Barda, V. Scaiewicz, M. Mills, T. Mueller, A. Gonzalez-Rodriguez, A. M. Valverde, T. Iwawaki, Y. Nahmias, R. Xavier, R. T. Chung, B. Tirosh, and O. Shibolet. 2013. ’CHOP is a critical regulator of acetaminophen-induced hepatotoxicity’, J Hepatol, 59: 495–503.

65. Walsh, K., H. Katow, H. Junn, D. Vasudevan, C. Dieterich, and H. D. Ryoo. 2025. ’4EHP and NELF-E regulate physiological ATF4 induction and proteostasis in disease models of Drosophila’, Nat Commun, 17: 626.

66. Wu, J., D. T. Rutkowski, M. Dubois, J. Swathirajan, T. Saunders, J. Wang, B. Song, G. D. Yau, and R. J. Kaufman. 2007. ’ATF6alpha optimizes long-term endoplasmic reticulum function to protect cells from chronic stress’, Dev Cell, 13: 351–64.

67. Yamaguchi, H., and H.G. Wang. 2004. ’CHOP is involved in endoplasmic reticulum stress-induced apoptosis by enhancing DR5 expression in human carcinoma cells’, J Biol Chem, 279: 45495–502.

68. Yamamoto, K., T. Sato, T. Matsui, M. Sato, T. Okada, H. Yoshida, A. Harada, and K. Mori. 2007. ’Transcriptional induction of mammalian ER quality control proteins is mediated by single or combined action of ATF6alpha and XBP1’, Dev Cell, 13: 365–76.

69. Yang, C., Z. Wang, Y. Hu, S. Yang, F. Cheng, J. Rao, and X. Wang. 2022. ’Hyperglycemia-triggered ATF6-CHOP pathway aggravates acute inflammatory liver injury by beta-catenin signaling’, Cell Death Discov, 8: 115.

70. Yang, Y., L. Liu, I. Naik, Z. Braunstein, J. Zhong, and B. Ren. 2017. ’Transcription Factor C/EBP Homologous Protein in Health and Diseases’, Front Immunol, 8: 1612.

71. Ye, B., M. Yue, H. Chen, C. Sun, Y. Shao, Q. Jin, C. Zhang, and G. Yu. 2024. ’YAP/TAZ as master regulators in liver regeneration and disease: insights into mechanisms and therapeutic targets’, Mol Biol Rep, 52: 78.

72. Yu, S., S. Meng, M. Xiang, and H. Ma. 2021. ’Phosphoenolpyruvate carboxykinase in cell metabolism: Roles and mechanisms beyond gluconeogenesis’, Mol Metab, 53: 101257.

73. Zhang, K., S. Wang, J. Malhotra, J. R. Hassler, S. H. Back, G. Wang, L. Chang, W. Xu, H. Miao, R. Leonardi, Y. E. Chen, S. Jackowski, and R. J. Kaufman. 2011. ’The unfolded protein response transducer IRE1alpha prevents ER stress-induced hepatic steatosis’, EMBO J, 30: 1357–75.

74. Zhang, P., J. Iwasaki-Arai, H. Iwasaki, M. L. Fenyus, T. Dayaram, B. M. Owens, H. Shigematsu, E. Levantini, C. S. Huettner, J. A. Lekstrom-Himes, K. Akashi, and D. G. Tenen. 2004. ’Enhancement of hematopoietic stem cell repopulating capacity and self-renewal in the absence of the transcription factor C/EBP alpha’, Immunity, 21: 853–63.

75. Zhang, P., and K. Reue. 2017. ’Lipin proteins and glycerolipid metabolism: Roles at the ER membrane and beyond’, Biochim Biophys Acta Biomembr, 1859: 1583–95.

76. Zhou, S., Z. Rao, Y. Xia, Q. Wang, Z. Liu, P. Wang, F. Cheng, and H. Zhou. 2023. ’CCAAT/Enhancer-binding Protein Homologous Protein Promotes ROS-mediated Liver Ischemia and Reperfusion Injury by Inhibiting Mitophagy in Hepatocytes’, Transplantation, 107: 129–39.

77. Zinszner, H., M. Kuroda, X. Wang, N. Batchvarova, R. T. Lightfoot, H. Remotti, J. L. Stevens, and D. Ron. 1998. ’CHOP is implicated in programmed cell death in response to impaired function of the endoplasmic reticulum’, Genes Dev, 12: 982–95.

