## Supplemental Figures S1-S6 for "CHOP promotes the transition to chronic integrated stress response signaling with suppression of hepatocyte identity"

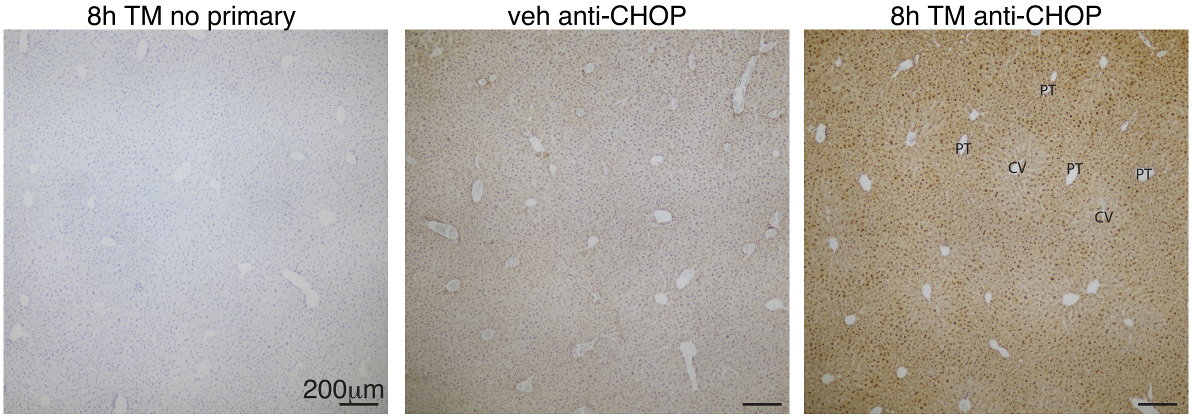


**Figure S1**, Related to Figure 1
Wild-type animals were injected with vehicle (veh) or TM for 8h, followed by hematoxylin counterstain and immunohistochemistry to detect CHOP, or without primary antibody as a control (left panel). Portal triad (PT) and central vein (CV) are indicated, showing more intense nuclear CHOP staining in periportal hepatocytes.

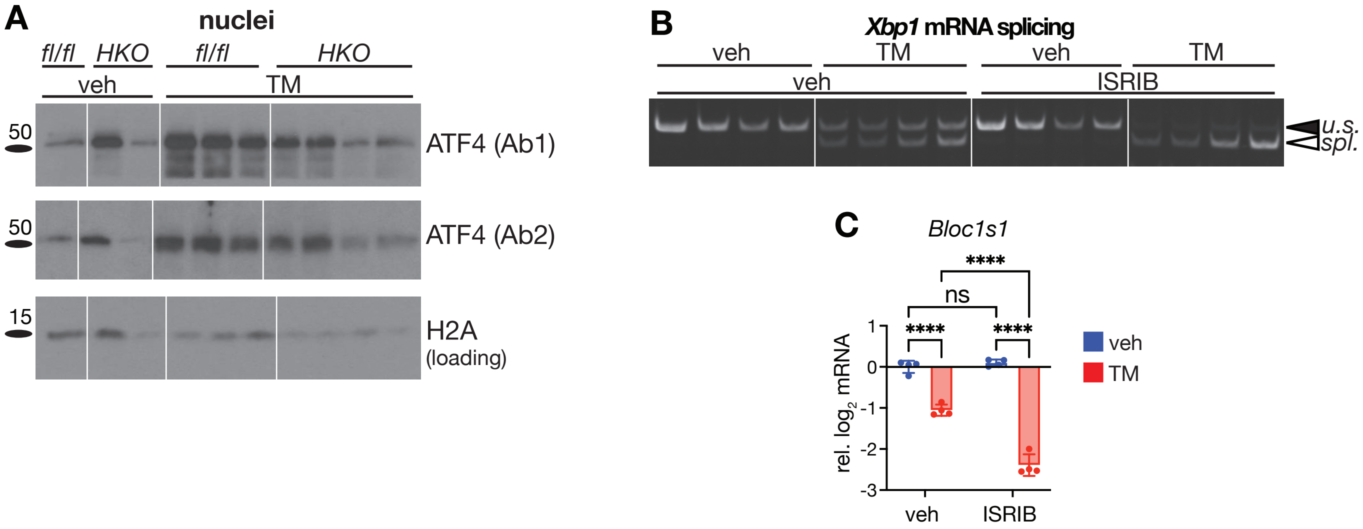


**Figure S2**, Related to Figure 2

**(A)** Nuclei were purified by centrifugation through a cushion of 2.3M sucrose from *Chop*$^{fl/fl}$ or *Chop*$^{HKO}$ animals treated with vehicle or TM as indicated. Two separate antibodies were used to detect ATF4. **(B, C)** Mouse primary hepatocytes were treated for 8h with or without 1 µg/ml TM and 1 µM ISRIB as indicated. IRE1 activation was detected by *Xbp1* mRNA splicing (B) or suppression of mRNA of the RIDD target *Bloc1s1* (C).

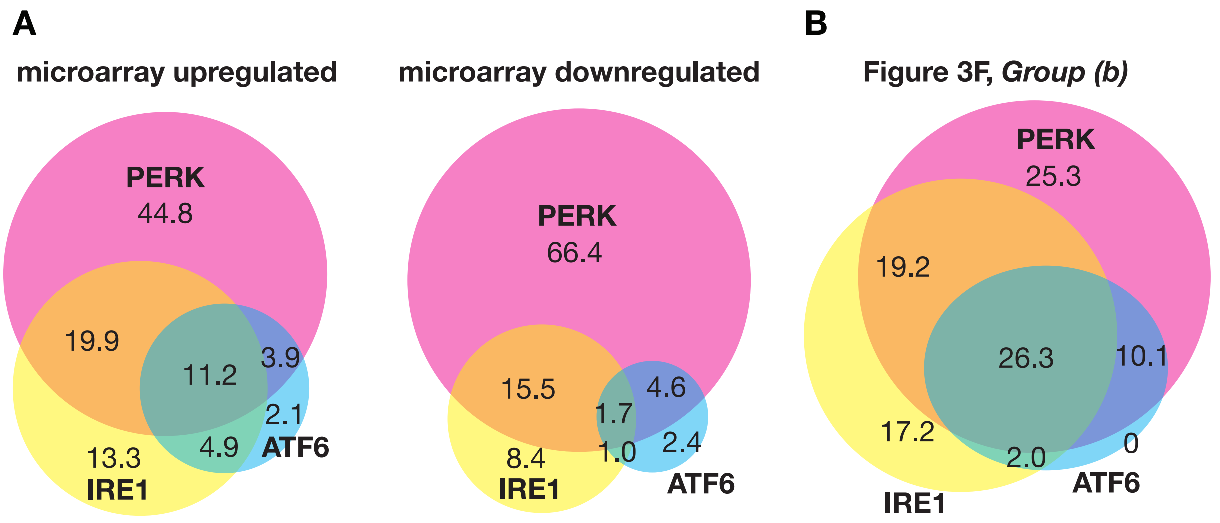


**Figure S3**, Related to Figure 3

**(A)** Data were aggregated from RNA microarrays taken from the livers of animals challenged with 1 mg/kg TM for 8h in *Perk*$^{LKO}$ (Teske et al., 2011), *Ire1*$\alpha^{LKO}$ (Zhang et al., 2011), or *Atf6*$\alpha^{-/-}$ (Rutkowski et al., 2008) mice. Probesets significantly upregulated or downregulated by TM treatment in matching wild-type animals from all three experiments were then assessed for whether that regulation was significantly diminished by deletion of one or more UPR sensor. Percentages in each group are shown (because of rounding, percentages might not sum to 100). Genes that could not be linked to one or more of the three pathways are not shown. **(B)** The genes in group (d) of Figure 3F were assessed in the same microarrays, in the same manner as in (A). For genes represented by more than one probeset, average regulation across all probesets was used. It can be seen that a substantially greater percentage of genes upregulated by the UPR and resistant to loss of CHOP are those that are also regulated by IRE1 and/or ATF6 compared to the overall pool of regulated genes

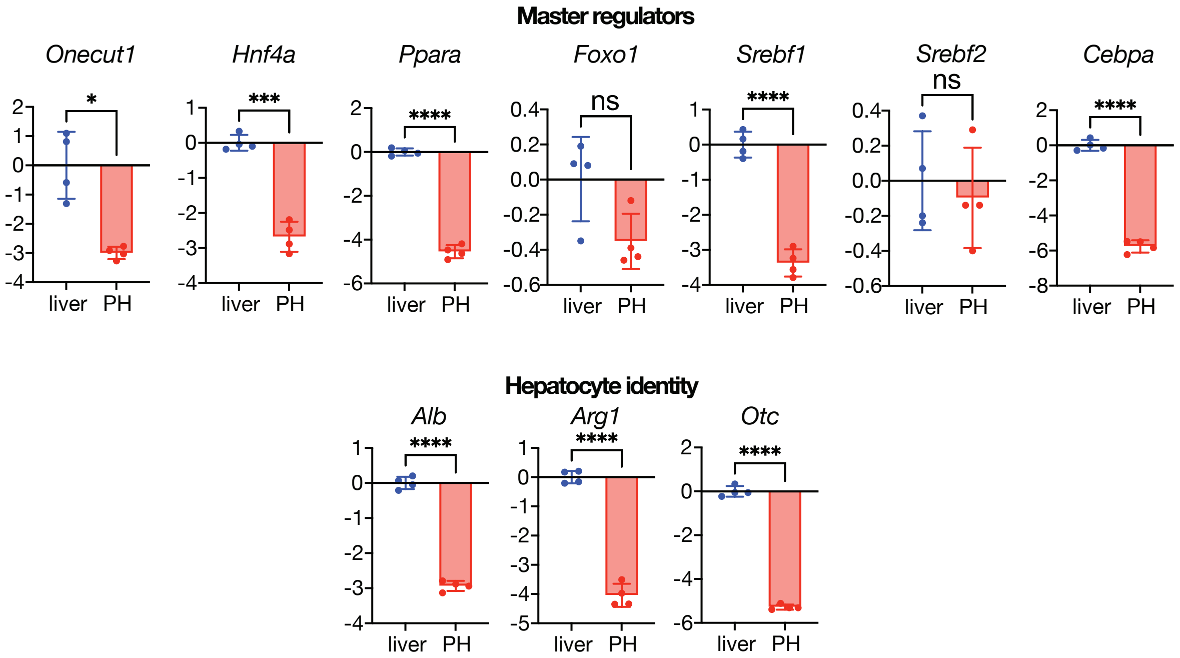


**Figure S4**, Related to Figure 5
Expression of the indicated genes was compared in primary hepatocytes isolated and allowed to rest for one day after isolation (as in Figure 5) and in liver tissue. Selected transcriptional master regulators of hepatocyte identity are shown above, and hepatocyte marker genes below. Statistical analysis by unpaired two-tailed t-test.

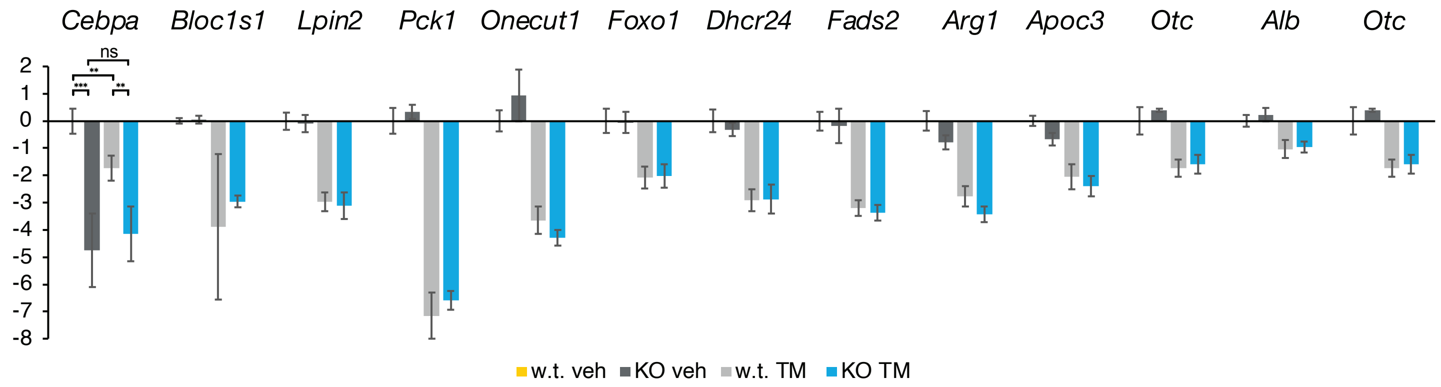


**Figure S5**, Related to Discussion
*Cebpa*$^{fl/fl}$ animals were treated with AAV-TBG-GFP (w.t.) or AAV-TBG-CRE (KO), and then two weeks later challenged with 1 mg/kg TM or vehicle for 8h. Expression of the indicated genes is shown. Other than *Cebpa* itself, none of the genes tested was significantly different between genotypes. n = 3-5 animals per group.

Figure 1B

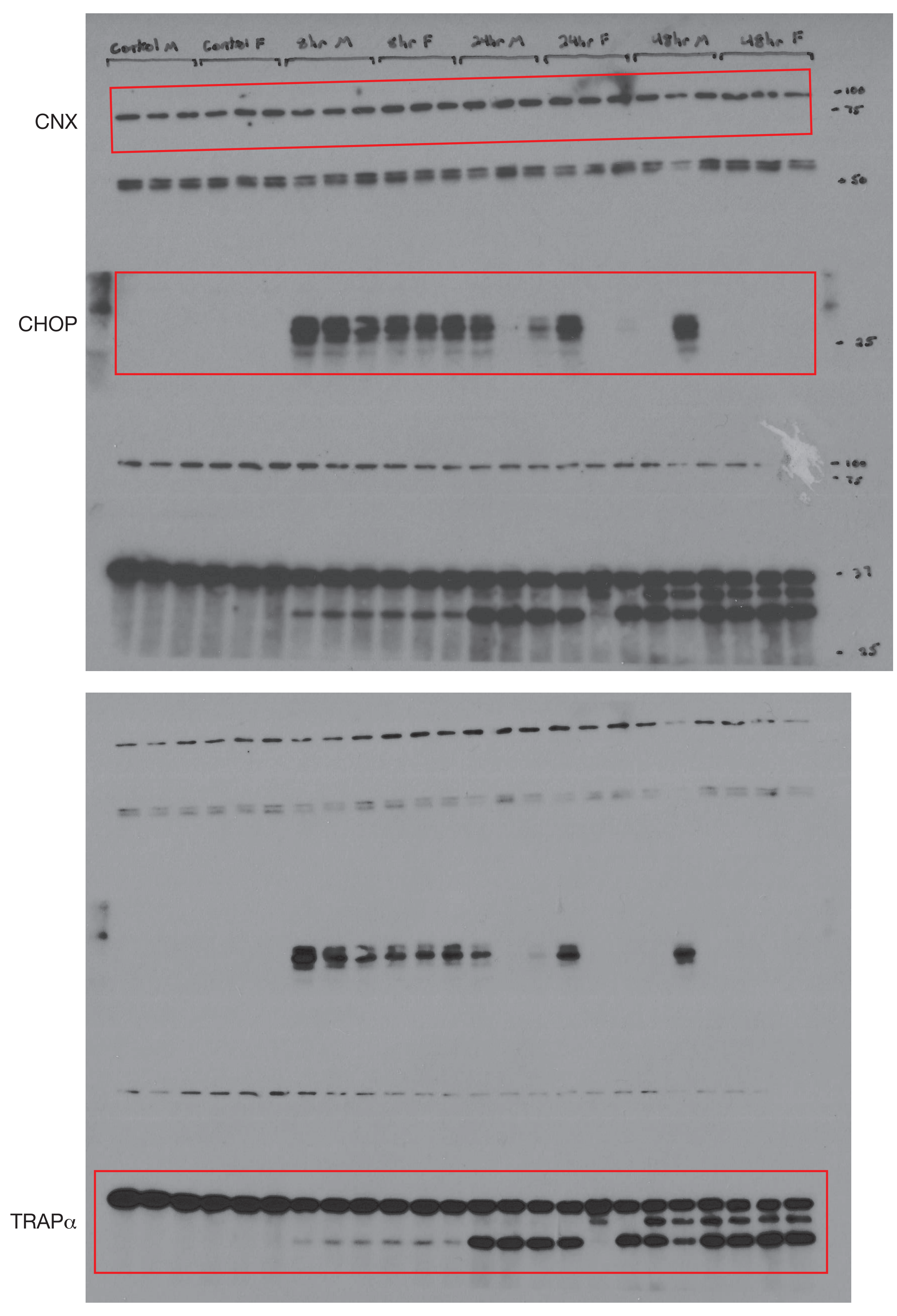


Figure 2F

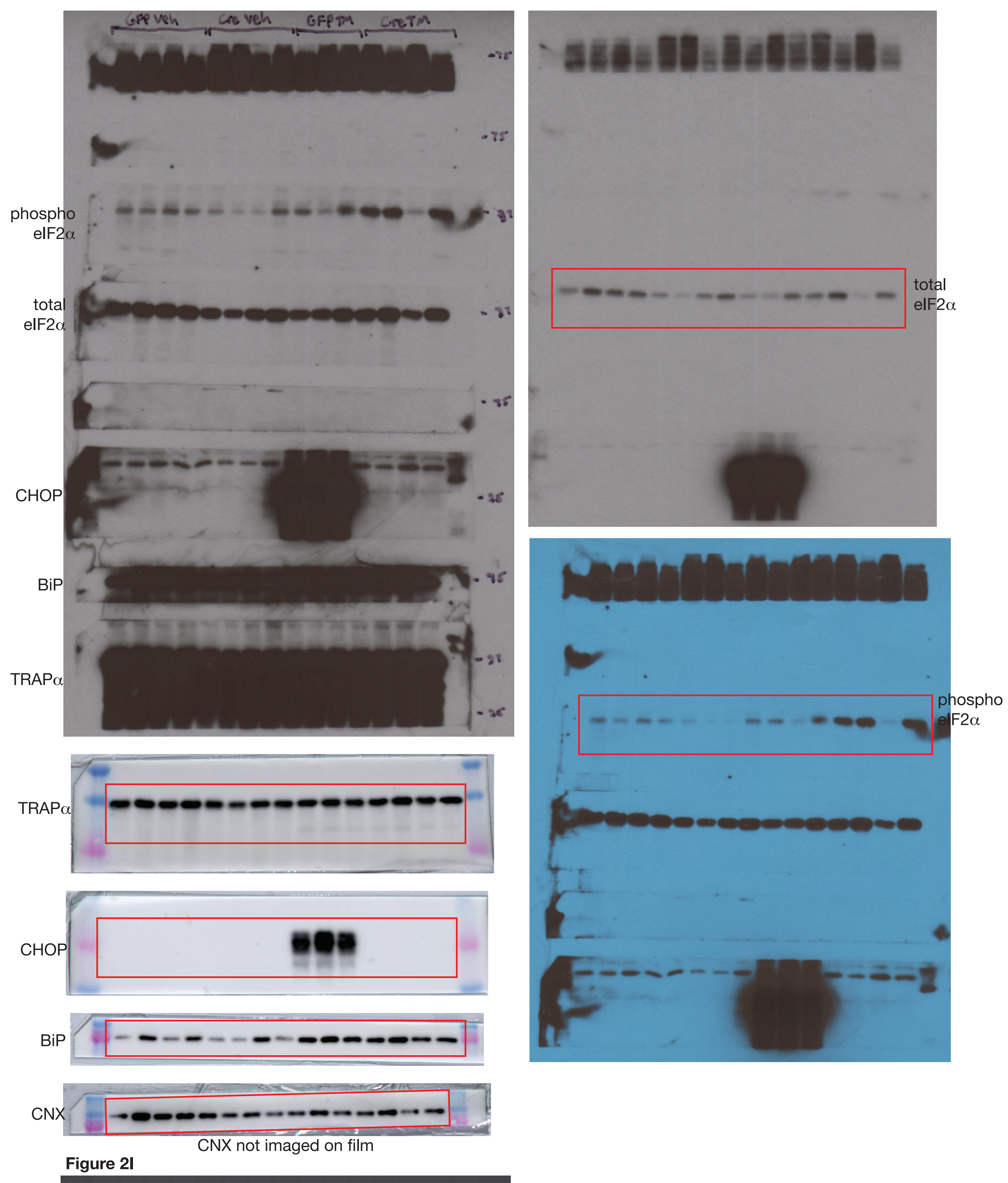


Figure 2I

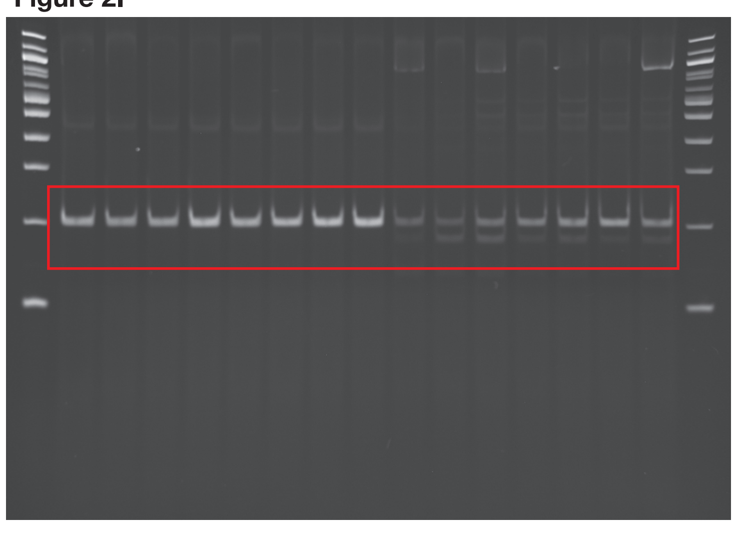

Figure S2A

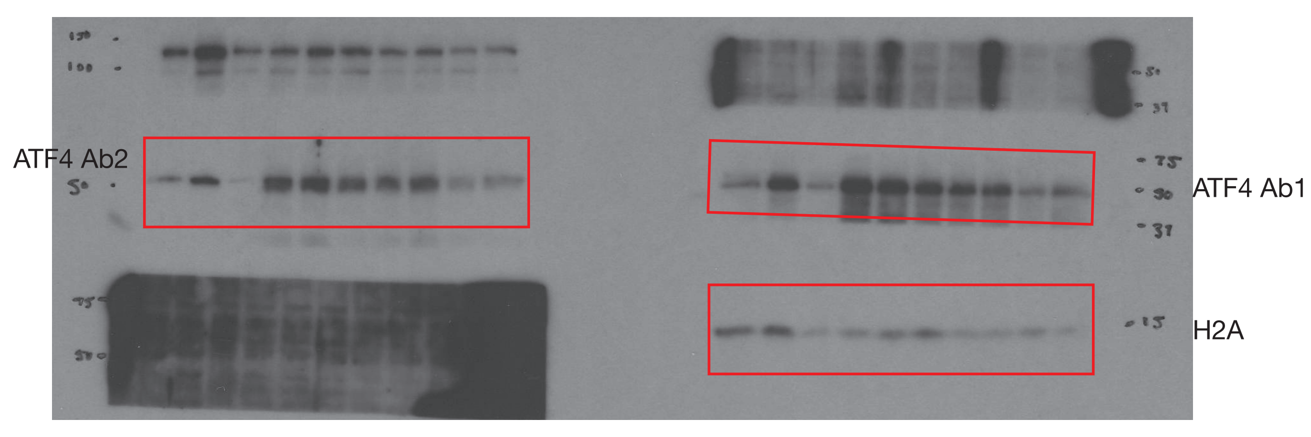


Figure S2B

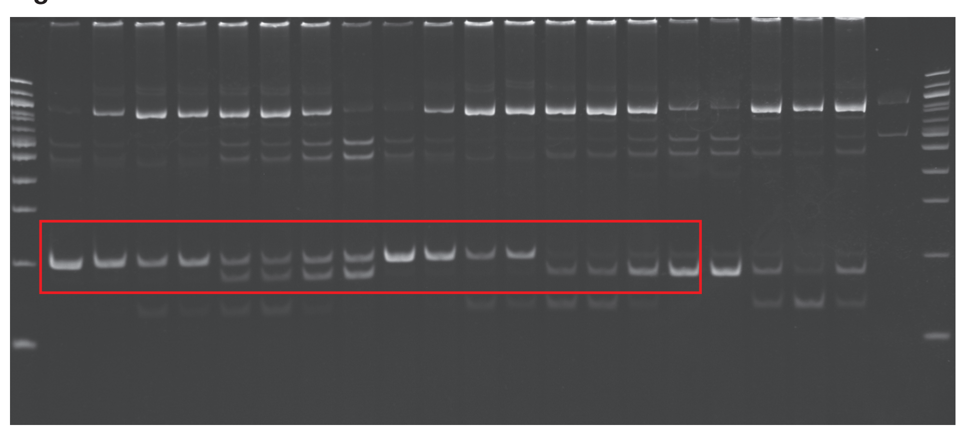

Figure 3B

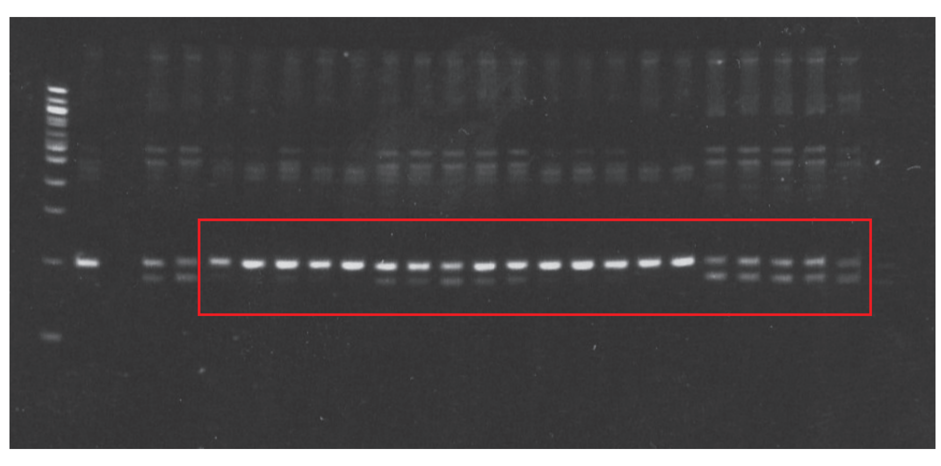

Figure 6B

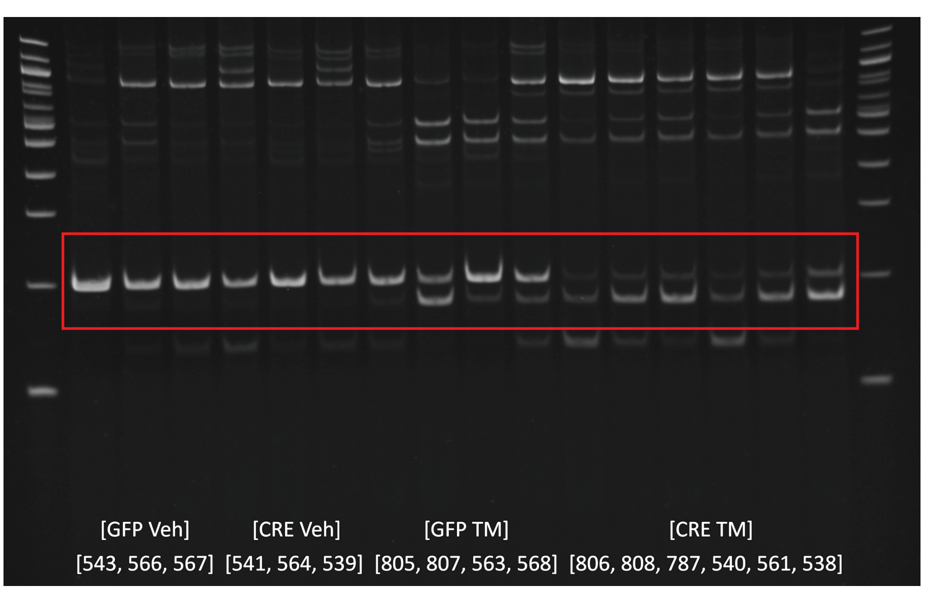


Figure 6E

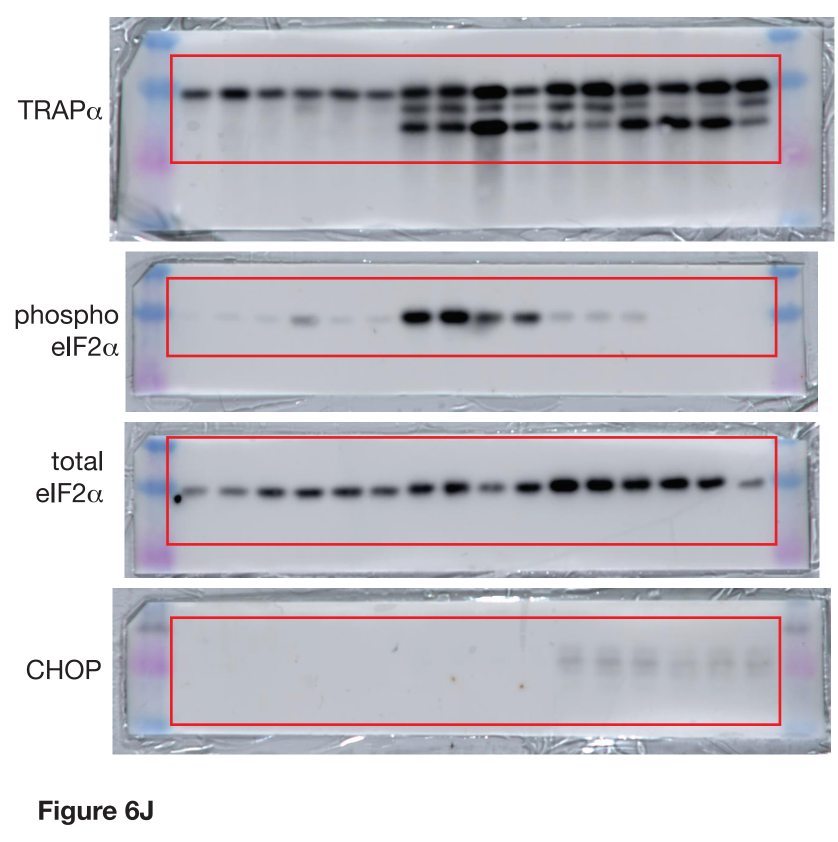


Figure 6J

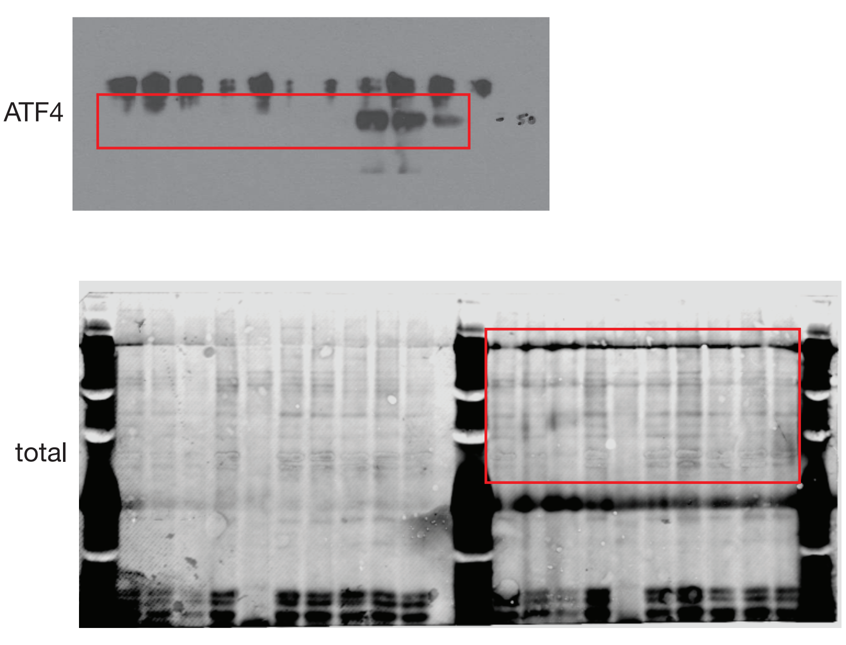
